# Regionalized cell and gene signatures govern oesophageal epithelial homeostasis

**DOI:** 10.1101/2024.02.21.581361

**Authors:** David Grommisch, Harald Lund, Evelien Eenjes, Anais Julien, Christian Göritz, Robert A. Harris, Rickard Sandberg, Michael Hagemann-Jensen, Maria Genander

## Abstract

Regionalized disease prevalence is a common feature of the gastrointestinal tract. Herein, we employed regionally resolved Smart-seq3 single-cell sequencing, generating a comprehensive cell atlas of the adult mouse oesophagus. Characterizing the oesophageal axis, we unveil non-uniform distribution of epithelial basal cells, fibroblasts and immune cells. In addition, we reveal a position-dependent, but cell subpopulation-independent, transcriptional signature, collectively generating a regionalized oesophageal landscape.

Combining *in vivo* models with organoid co-cultures, we demonstrate that proximal and distal basal progenitor cell states are functionally distinct. We find that proximal fibroblasts are more permissive for organoid growth compared to distal fibroblasts and that the immune cell profile is regionalized in two dimensions, where proximal-distal and epithelial-stromal gradients impact epithelial maintenance. Finally, we predict and verify how WNT-, BMP-, IGF-and NRG-signalling are differentially engaged along the oesophageal axis.

We establish a cellular and transcriptional framework for understanding oesophageal regionalization, providing a functional basis for epithelial disease susceptibility.

## Introduction

The gastrointestinal tract is an epithelial-lined muscular tube extending from the oral cavity to the anus. Although the general architecture of the tract is conserved from beginning to end, the presence of unique epithelial structures and cell types enables each organ to perform distinct digestive functions. Characterization of the adult gastrointestinal epithelium demonstrates that developmentally established gene expression patterns are retained, and genetic ablation of regionally restricted epithelial transcription factors changes adult cell differentiation, tissue architecture and function ^1–4^. Similarly, regionally organized epithelial expression of innate immune signalling components determines location-specific responses to pathogens ^5^, and activation of tumour promoters link region-specific differences in epithelial cell states to oncogenic pathway activity ^6^. Taken together this indicates that regional epithelial gene expression profiles impact mechanisms of adult epithelial homeostasis as well as disease.

Stem and progenitor cells commonly localise to distinct anatomical niches, enabling exclusive interaction with non-epithelial cell types and access to molecular determinants, collectively acting to maintain stemness and prevent differentiation ^7–12^. Characterization of stem cell niches has identified mechanisms driving gastrointestinal regionalization. Regional comparison of intestinal fibroblast subpopulations using single-cell sequencing reveals identical subpopulation clusters and frequencies when comparing the small intestine to the colon ^13^. However, stromal expression of WNT-modulating genes as well as the distance between intestinal stem cells and BMP-producing fibroblasts varies along the proximal-distal axis, leading to an overall higher WNT-activity in the colonic epithelium compared to in the small intestine. This work exemplifies how differences in niche architecture combined with locally defined gene expression profiles govern local stem cell behaviours and regionalized mechanisms of epithelial maintenance ^13^.

In contrast to intestine, the oesophageal epithelium lacks crypts and no anatomically defined progenitor niches have been described. Interestingly, however, human adeno- and squamous cell carcinomas and inflammatory pathologies predominantly target the lower two thirds of the oesophagus ^14–17^. While differences in muscle layers, innervation, and distribution of immune cell populations along the human oesophageal axis are described ^18,19^, the relevance of these cellular differences are currently unknown. We have recently reported that a subset of functionally distinct, *Troy*-positive, oesophageal progenitor cells localises to discrete epithelial regions in a proximal-distal gradient ^20^. *Troy*-expressing progenitors are less proliferative than their *Troy*-negative neighbours, only to become preferentially activated upon regenerative challenge. This suggests that regionally defined cellular and transcriptional profiles could govern local progenitor cell states and disease susceptibility also in the oesophagus.

Herein, we employed regionally resolved Smart-seq3 single-cell sequencing ^21,22^ to characterize the adult mouse oesophageal proximal-distal axis. We capture all major cell populations in the oesophagus, identify and compare proximal-to-distal progenitor-niche networks and molecular gradients and combine *in vivo* models with organoid co-cultures ^23^ to functionally demonstrate proof-of-concept cellular interactions and signalling dependencies. This study provides a comprehensive and importantly, regionally resolved, oesophageal cell atlas, challenging our current understanding of oesophageal tissue homeostasis and providing insights into clinically relevant disease susceptibility.

## Results

### Regionally resolved single-cell transcriptomics

To begin to understand regionalization of the mouse oesophageal tube, we performed H&E staining comparing proximal to distal oesophageal cross-sections allowing for a macroscopic comparison of the epithelium, stroma and the muscularis externa. We found changes in the oesophageal architecture along the axis (Figure 1A), where a distal thickening of the oesophageal tube was largely driven by an expansion of the muscularis externa. To understand if proximal-distal differences extended to the epithelium and stroma we employed single-cell transcriptomics. We divided the oesophagus into three parts (proximal, middle, and distal), mechanically removed the muscularis externa and enzymatically separated the epithelium from the stroma ^23^. Single cells were FACS-isolated and Smart-seq3 transcriptomes from the six tissue fractions were collected (Figure 1B) ^21,22^. Following quality control (see methods section) a total of around 14,162 cells passed the selected criteria and were subjected to downstream analysis.

**Figure 1.**
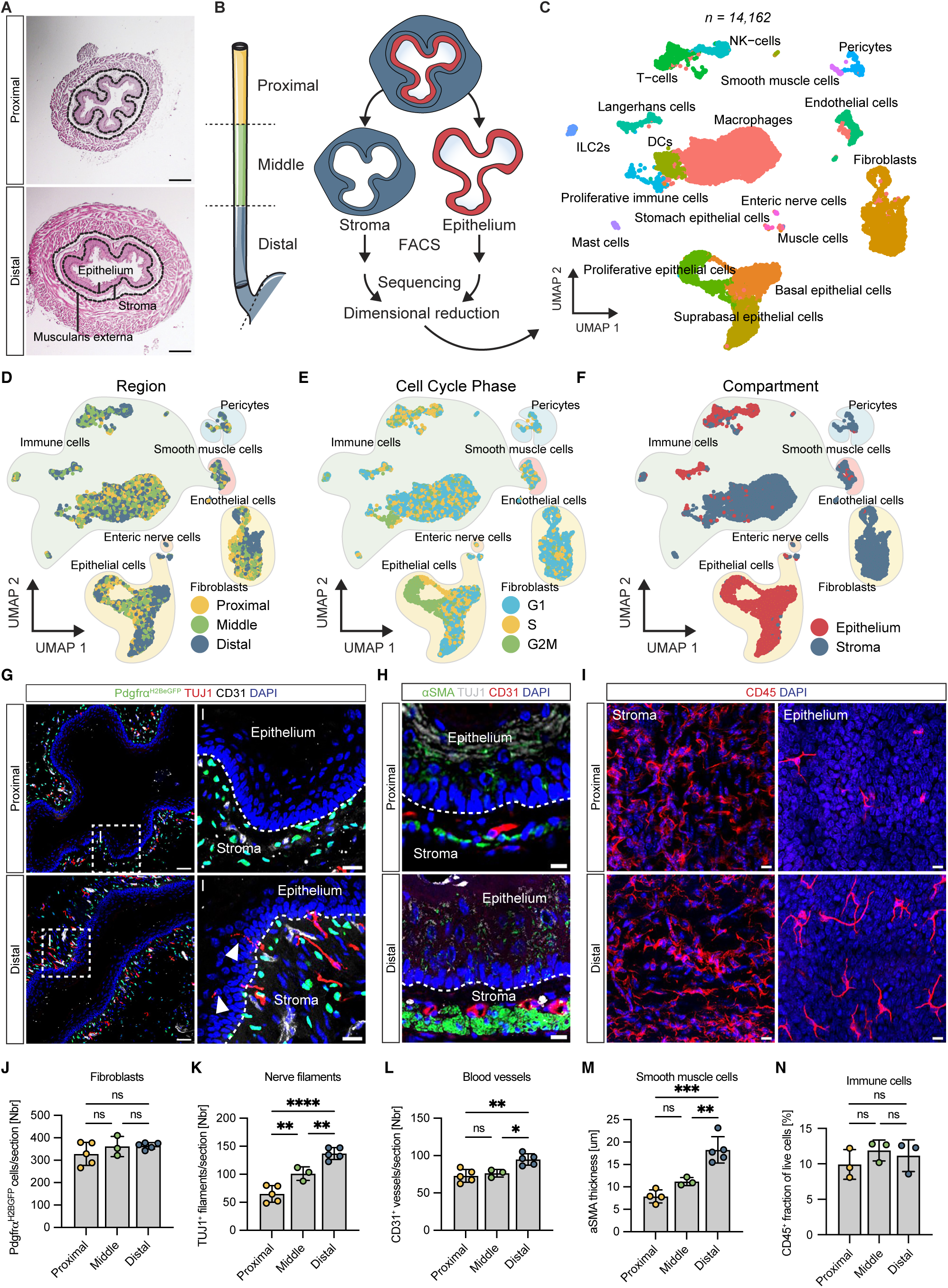
Regionally resolved single-cell transcriptomics. (A) Haematoxylin-Eosin staining of cross sections of the proximal and distal oesophagus. Dotted lines demark borders between epithelium, stroma, and muscularis externa. Scale bar: 200µm. (B) Illustration of proximal-middle-distal tissue division and epithelial-stromal separation for single-cell RNA sequencing. (C) Uniform manifold approximation and projection (UMAP) dimensional reduction of all sequenced oesophageal cells. (D) UMAP colour-coded by region-of-origin. (E) UMAP colour-coded by inferred cell-cycle stage. (F) UMAP colour-coded by oesophageal epithelial-stromal compartment. (G) Immunofluorescent staining of TUJ1 (red, nerve fibres), CD31 (white, vessels), and Pdgfrα^H2BeGFP^ (green, fibroblasts) on cross-sections. Scale bar = 50µm and inlets 20µm. (H) Immunofluorescent staining of αSMA (green, muscularis externa), TUJ1 (white), and CD31 (red) of cross-sections. Scale bar = 20µm. (I) Immunofluorescent localization of CD45-positive immune cells (red) in stromal and epithelial whole mounts, respectively. Scale bar = 10µm. (J) Quantification of Pdgfrα^H2BeGFP^-positive nuclei in cross-sections. (K) TUJ1+ nerve fibres abundance on cross-sections. (L) CD31+ vessel abundance on cross-sections. (M) Thickness of the αSMA-positive muscularis externa on oesophageal cross-sections. (N) Flow cytometry-based quantification of CD45-positive immune cells in proximal, middle, and distal oesophageal segments. n = 3 (2-3 mice per n). (J, K, L, M, N) Ordinary One-way ANOVA corrected for multiple comparisons (Tukey). n = 3-5 mice. ns p > 0.05, * p < 0.05, ** p < 0.01, *** p < 0.001. Dotted lines mark the epithelial-stromal border.

Unsupervised hierarchical clustering identified 18 different cell types and states (Figure 1C and S1A) which were further grouped into seven major cell populations (epithelial, fibroblast, immune cells, pericytes, smooth muscle cells, enteric nerve cells, and endothelial cells) based on known gene expression of these cell types (Figure S1B and S1C). We then visualized cells based on their region-of-origin. While all cell type clusters were represented in all regions, we noted enrichment of cells derived from either the proximal and distal oesophagus to specific parts of the UMAP (most prominently evident for epithelial, fibroblasts and immune cells) (Figure 1D). Inferring cell cycle stages revealed that epithelial cells clustered based on their proliferation dynamics, whereas fibroblasts were predominantly present in G1 and clustered independently of the cell cycle (Figure 1E). Finally, we confirmed that oesophageal cells segregated based on their epithelial-stromal origin (Figure 1F), but that immune cells were identified within both the epithelial and stromal fractions, likely representing discrete immune cell types (Figure 1F). Collectively, this dataset represents a comprehensive, regionally resolved cell atlas indicating the existence of region-specific subpopulations and/or a non-uniform distribution of key cell populations along the oesophageal axis.

### Characterization of regional stromal cell types

To understand if the regional cell clustering (Figure 1D) could represent local microenvironments, or progenitor niches, we quantified the number of *Pdgfrα*^H2BeGFP^ reporter positive fibroblasts in the stroma (Figure 1G and 1J) but found no significant differences in the total number of fibroblasts when comparing the proximal, middle and distal oesophagus. Whereas the frequency of TUJ1-positive peripheral nerve endings was significantly enriched (Figure 1G-H, 1K and S1D-E, S1G-I) (Supplemental Table 1 and 2) and nerve endings extended into the epithelium in the distal compared to the middle and proximal oesophagus (Figure 1G, white arrows in inset), we only found a modest increase in the number of CD31-positive vessels in the distal oesophagus (Figure 1G-H,1L and S1D, S1F-I). In contrast, the thickness of the αSMA-positive muscularis externa increased significantly distally (Figure 1H, 1M and S1G-I). Immunohistochemistry suggested, and flow cytometry quantification confirmed an even distribution of CD45-positive immune cells within the oesophageal stroma (Figure 11 and 1N), but we detected an enrichment of intraepithelial immune cells in the distal compared to the proximal epithelium (Figure 1I). These data demonstrate that several stromal cell types known to impact epithelial progenitor behaviours are unevenly distributed along the oesophageal axis ^24,25^. In addition, the even distribution of both stromal fibroblasts and immune cells indicates that further in-depth characterization of PDGFRα- and CD45-positive subtypes is required to fully understand the proximal-distal cellular architecture (Figure 1D).

### Regional gene expression signatures define the oesophageal epithelium

To understand the changes along the oesophageal axis in more detail, we subclustered and re-analysed all epithelial cells. We confirmed that epithelial cells derived from the epithelial cell fraction (Figure S2A) and identified 12 epithelial subpopulations (Figure 2A) (Supplemental Table 3). In addition to four subpopulations of cycling basal cells (Proliferative 1-4) and three subpopulations undergoing cell state transitions associated with oesophageal differentiation (Suprabasal 1-3) we identified five basal subpopulations (Basal 1-5) (Figure 2A and S2B-D). Some basal subpopulations located predominantly to the proximal and middle oesophagus (Basal 1 and Basal 2), whereas others were mainly located in the distal epithelium (Basal 3) (Figure 2A-C and S2E). Gene Ontology analysis comparing the five basal clusters revealed functional differences, encompassing extracellular matrix organization (Basal 1), tissue development (Basal 2) WNT and planar cell polarity signalling (Basal 3), mRNA processing (Basal 4) and ribosomal biogenesis and response to viral infection (Basal 5) (Supplemental Table 4) ^26^. These data suggest that subpopulations of basal cells exist and raises the possibility that cell location infers basal progenitor cell states, or conversely, that cell states mediate tissue regionality.

**Figure 2.**
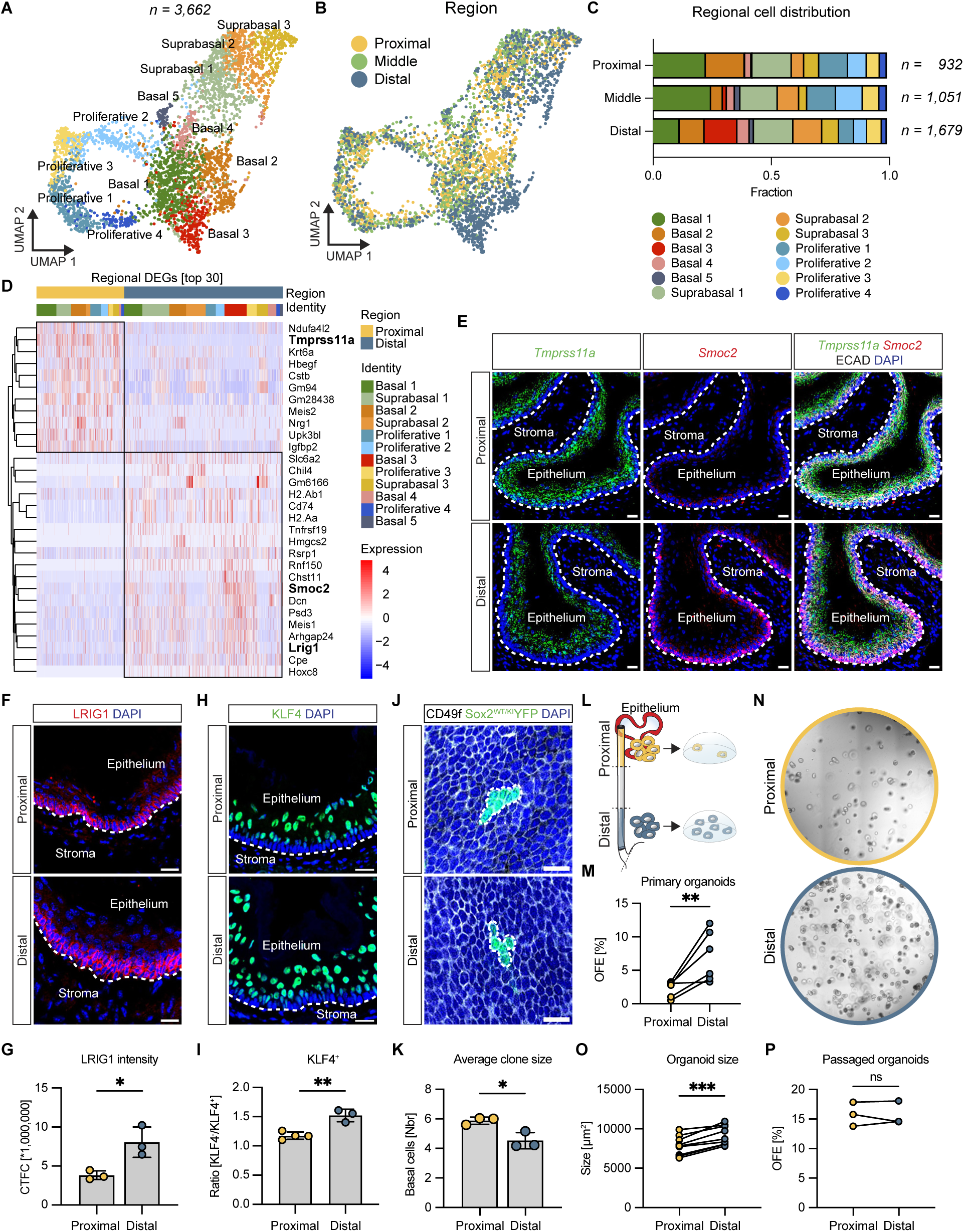
Regional gene expression signatures define the oesophageal epithelium. (A) UMAP of subclustered epithelial cell identities. (B) UMAP of epithelial cells colour-coded by oesophageal region-of-origin. (C) Quantification of the distribution of epithelial subpopulations within each sequenced proximal-distal region. Cell numbers are normalised within respective region, displaying fractions. (D) Heatmap of top 30 differentially expressed genes comparing all proximal and distal epithelial cells. (E) *In situ* hybridization of *Tmprss11a* (green) and *Smoc2* (red) on cross-sections of the proximal and distal oesophagus. Sections are counterstained with E-CADHERIN (ECAD, white) and DAPI (blue). Scale bar = 20µm. (F) LRIG1 (red) staining of the proximal and distal oesophagus. Sections are counterstained with DAPI. Scale bar = 20µm. (G) Quantification of corrected total cell fluorescence (CTCF) of LRIG1 in cross-sections of the proximal and distal oesophagus. (H) KLF4 (green) staining of the proximal and distal oesophagus. Scale bar = 20µm. (I) Quantification of KLF4-negative (basal) to KLF4-positive (suprabasal) cells comparing proximal and distal cross-sections. (J) Whole mount images of Sox2^CreER^:YFP-positive basal clones after 3 months of tracing in proximal and distal oesophagus. Wholemounts were counterstained with CD49f (white) and DAPI (blue). Scale bar = 20µm. (K) Quantification of average basal cell number within Sox2^CreER^:YFP-positive clones. (L) Schematic illustration of organoid formation strategy, isolating basal progenitor cells from the proximal (yellow) or distal (blue) epithelium. (M) Organoid forming efficiency (OFE) at day 6 of organoid culture comparing proximal and distal OFE in cultures established from the same oesophagus. n=6. (N) Representative brightfield images of epithelial oesophageal organoids derived from the proximal and distal oesophageal epithelium, respectively. Scale bar = 100µm. (O) Quantification of the size of primary proximal and distal organoids. Proximal and distal organoids established from the same oesophagus are compared. n=9. (P) Organoid forming efficiency of proximal and distal derived organoids after passaging the same number of FACS isolated organoid cells. n=3. (G, I, K, M) Two-sided unpaired student’s t-test. (M, O, P) Two-sided paired student’s t-test. ns p > 0.05, * p < 0.05, ** p < 0.01, *** p < 0.001. Dotted lines mark the epithelial-stromal border.

To uncouple regionality from cell state, we compared the transcriptional profiles of all epithelial cells isolated from the proximal and distal epithelia (Figure 2D) (Supplemental Table 5). Genes identified as proximal (*Tmprss11a*) or distal (*Smoc2* and *Lrig1*) signature genes were also found differentially expressed between proximal and distal cells from several subclusters (Figure 2D and S2F) indicating that regional transcriptional signatures across epithelial subclusters could impact basal progenitor cell functionality.

*In situ* hybridization confirmed prominent *Tmprss11a* mRNA expression in proximal basal and suprabasal epithelial cells (Figure 2E). In the distal epithelium, *Tmprss11a* mRNA expression was lower and predominantly localised to suprabasal cells (Figure 2E). Conversely, *Smoc2* mRNA and LRIG1 protein expression were low in the proximal epithelium but increased in basal progenitor cells of the distal epithelium (Figure 2E-G). In addition, LRIG1 immunoreactivity extended into the first suprabasal layer in the distal, but not the proximal epithelium (Figure 2F). Taken together, this work suggests that both basal and suprabasal oesophageal epithelial cells are affected by their location along the proximal-distal oesophageal axis.

## Functional basal progenitor states are regionally specified

We then addressed if regional gene expression signatures corresponded to distinct mechanisms of tissue maintenance. Using KLF4 to label differentiated suprabasal cells ^27^, we quantified the ratio of basal to suprabasal cells in proximal and distal epithelia respectively, revealing a significant relative increase in suprabasal cells in the distal epithelium (Figure 2H-I). To understand if differences in epithelial cell architecture corresponded to regional differences in basal progenitor cell behaviour, we administered a single dose of tamoxifen to Sox2^WT/KI^ (*Sox2-CreER^T2^:YFP*) mice ^28^ to induce recombination in random, single basal SOX2-positive progenitor cells ^20^. Quantification of resulting basal clone sizes at 3 months of tracing revealed that the average distal basal clone size was significantly smaller than the average proximal clone size (Figure 2J-K), suggesting that regional basal progenitor cell behaviour or dynamics is distinct, likely reflecting local fine-tuning of *in vivo* tissue maintenance.

To further understand regional basal progenitor cell behaviours, we compared the propensity of *in vitro* organoid formation of proximal and distal FACS isolated CD324/CD104 double positive basal progenitor cells. Strikingly, distal basal cells outperformed proximal basal cells in organoid formation efficiency (Figure 2L-N) and generated significantly larger organoids (Figure 2O). qPCR analysis from primary organoids confirmed that proximally enriched *Igfbp2* and *Tmprss11a* expression was maintained in culture (Figure S2G). To understand if regional differences in organoid forming capacity are determined by cell intrinsic factors or are influenced by local signalling cues, we passaged distal and proximal primary organoids and found that both regional differences in organoid forming capacity as well as gene expression signatures were lost or altered (Figure 2P and S2H-I). These data indicate that regional signalling cues impact functional basal progenitor states and behaviour.

## Identification of a distal basal progenitor subpopulation

We identified the Basal 3 cluster to be significantly enriched in the distal epithelium (Figure 2C). To characterize the identity of the Basal 3 subcluster, we determined differentially expressed genes in Basal 3 compared to all other basal cell subclusters (Supplemental Table 6). We found that Basal 3 cells expressed low levels of the proximal signature gene *Igfbp2* and elevated levels of *Pappa, Lrig1, Igfbp5,* and *Smoc2* (Figure 3A), all of which were enriched in the distal epithelium. Using the collective differential expression of *Pappa, Lrig1, Igfbp5,* and *Smoc2*, we confirmed faithful correlation with the Basal 3 subpopulation on the UMAP (Figure 3B-C and 2A).

**Figure 3.**
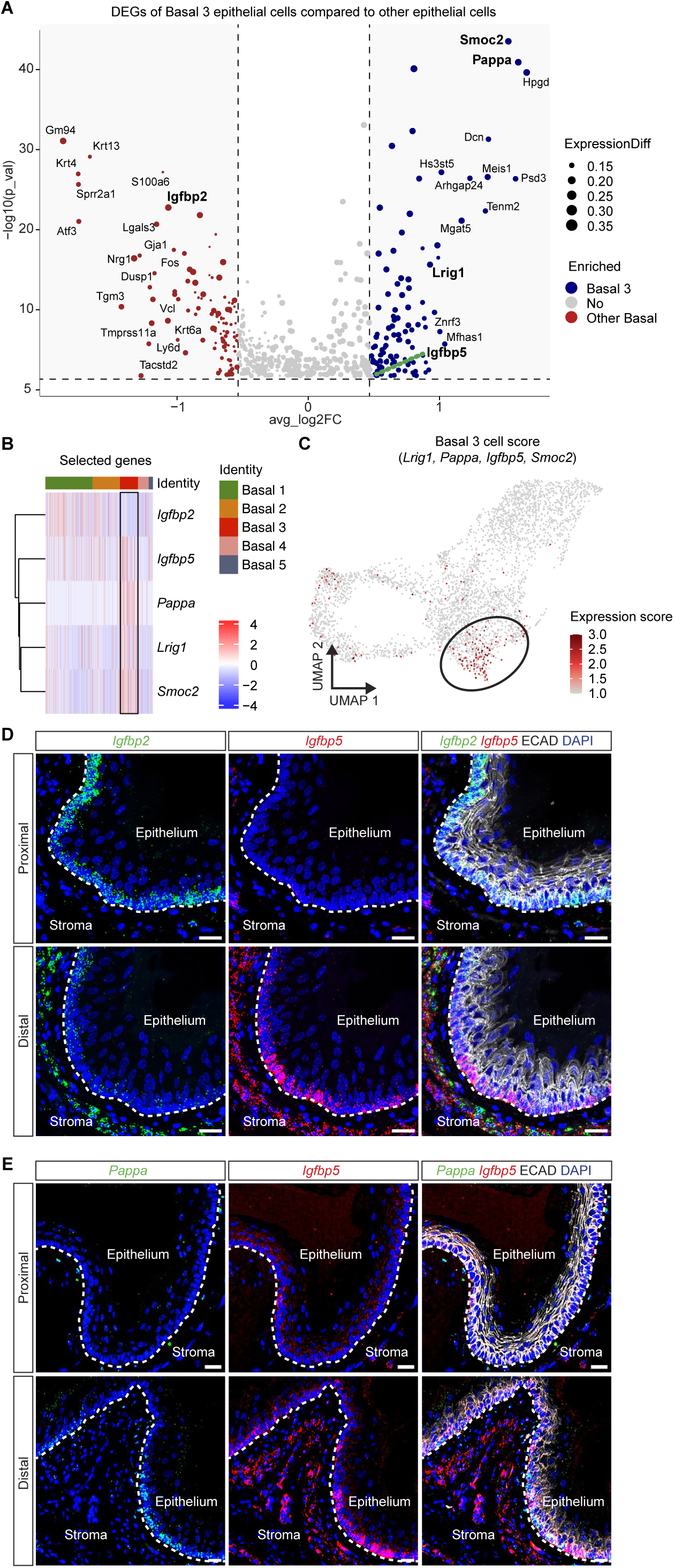
Identification of a distal basal progenitor subpopulation. (A) Volcano plot of differentially expressed genes between the Basal 3 subpopulation and other basal subpopulations (Basal 1, Basal 2, Basal 4 and Basal 5). Genes enriched in Basal 3 are showcased in blue, genes enriched in the other basal subpopulations are displayed in red. (B) Heatmap of selected DEGs comparing Basal 3 and other Basal subpopulations. (C) UMAP visualization of the Basal 3 gene expression score based on *Lrig1*, *Pappa*, *Igfbp5*, and *Smoc2* gene expression. (D) *In situ* hybridization for *Igfbp2* (green) and *Igfbp5* (red) as well as *Pappa* (green) and *Igfbp5* (red) (E) on proximal and distal oesophageal cross-sections. Sections are counterstained with E-CADHERIN (ECAD, white) and DAPI (blue). Scale bar = 20µm. Dotted line indicates epithelial-stromal border.

We then used *in situ* hybridization to localize Basal 3 progenitor cells. We confirmed decreasing proximal-to-distal expression of *Igfbp2,* whereas the opposite was true for *Igfbp5*, which displayed increased expression in the distal compared to the proximal epithelium (Figure 3D). In addition, *Igfbp5* was confined to discrete regions, or patches, of distal basal cells which expressed reduced levels of *Igfbp2*. Similarly to *Igfbp5*, *Pappa* expression was low in the proximal region only to increase in patches of distal basal cells epithelium. Dual labelling of *Pappa* and *Igfbp5* confirmed that the two genes were co-expressed in an alternating on-off pattern around the distal epithelial circumference (Figure 3E). These findings demonstrate the existence of a distal basal progenitor population which localizes to discrete regions of the epithelial circumference.

## Fibroblasts establish local signalling landscapes

Regional differences in organoid forming capacity were lost when primary organoids were passaged (Figure 2P), suggesting that environmental signalling cues, rather than cell intrinsic programs, impact basal progenitor cell states and behaviour. To address the contribution of fibroblasts to transcriptional and functional regionalization of the epithelium ^11,29,30^, we subclustered all previously annotated fibroblasts to unveil heterogeneity along the proximal-distal axis.

We identified and annotated eight fibroblast clusters (Figure 4A and S3A-C) (Supplemental Table 7) and uncovered functional differences linked to extracellular matrix production and organization (*Sfrp2*, *Anxa3* and *Spon1*), epithelial cell proliferation and differentiation (*Sfrp2*) and response to bacteria and virus (*Ifit3*) (Supplemental Table 8). In addition, we found that subclusters could be divided into previously described *Col15a1*^pos^ parenchymal (*Spon1*, *Kcnn3* and *Sfrp2*) and *Pi16*^pos^/*Dpp4*^pos^ adventitial (*Axna3* and *Ccl2*) fibroblasts (Supplemental Table 7), confirming the largely generalized cross-tissue identities of fibroblasts ^31^.

**Figure 4.**
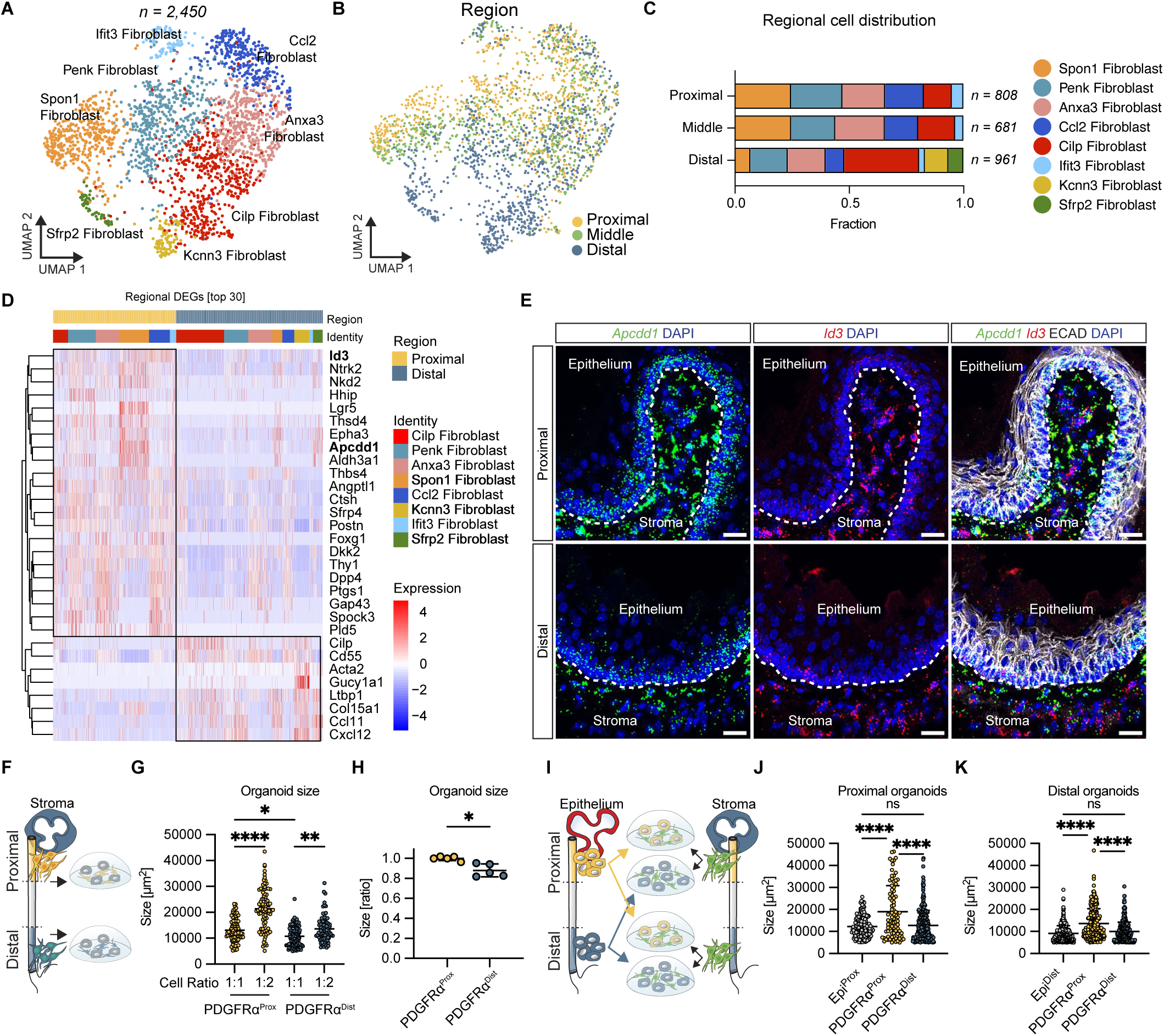
Fibroblasts establish local signalling landscapes. (A) UMAP of cell identities of subclustered *Pdgfrα*-positive fibroblasts. (B) UMAP of subclustered fibroblasts colour-coded by oesophageal region-of-origin. (C) Proximal-distal distribution of each fibroblast subpopulation. Cell numbers are normalised within respective region, displaying fractions. (D) Heatmap of top 30 DEGs comparing all proximal and distal fibroblasts. (E) *In situ* hybridization of *Apcdd1* (green) and *Id3* (red) on cross-sections of the proximal and distal oesophagus. Sections are counterstained with E-CADHERIN (ECAD, white) and DAPI (blue). Dotted line marks the epithelial-stromal border. Scale bar = 20µm. (F) Schematic illustration of organoid co-culture experiment. Regionally FACS isolated *Pdgfrα*^H2BeGFP^-positive fibroblasts were co-cultured with epithelial basal cells derived from the entire length of the epithelium. (G) Quantification of organoid size after co-culturing with regionally derived *Pdgfrα*^H2BeGFP^-positive cells at a 1:1 or 1:2 epithelial-to-fibroblast ratio. Increasing the number of fibroblasts positively affects organoid size. Each dot represents one organoid. (H) Quantification of relative average organoid size in 1:1 epithelial-to-fibroblast co-cultures, comparing the effect of fibroblasts originating from the proximal and distal stroma. Data is derived from 3 independent experiments and each dot represents a biological replicate paired from 2-3 mice. (I) Illustration of experimental setup pairing regionally derived oesophageal fibroblasts to regionally derived oesophageal epithelial cells. (J, K) Organoid size of regionally derived oesophageal *Pdgfrα*^H2BeGFP^-positive fibroblast co-cultured with proximally (J) or distally derived (K) epithelial basal progenitor cells. Each dot represents one organoid. (G, J, K) Ordinary One-way ANOVA corrected for multiple comparisons (Tukey). ns p > 0.05, * p < 0.05, ** p < 0.01, *** p < 0.001. (H) Two-sided unpaired student’s t-test. * p < 0.05, ** p < 0.01, *** p < 0.001, **** p < 0.0001. n = 3 – 5 (replicates are pooled from 2-5 mice).

Not all fibroblast subclusters were evenly located along the oesophageal axis (Figure S3D). Fibroblasts defined by their *Spon1*-expression were preferentially located in the proximal and middle oesophagus, whereas *Sfrp2*- and *Kcnn3*-positive fibroblasts were exclusively located to the distal oesophagus (Figure 4A-C and S3D). To understand if fibroblast regionality extended beyond the physical location of specific subpopulations, we compared the overall transcriptomes of proximal and distal fibroblasts (Supplemental Table 9). We identified *Apccd1*, *Lgr5*, *Sfrp4* and *Dkk2* (associated with WNT signalling) as well as *Hhip* (Hedgehog signalling effector) and *Id3* (BMP target gene) as being enriched, across several subclusters, in proximal fibroblasts (Figure 4D). In contrast, genes associated with TGFβ signalling (*Cilp* and *Ltbp1*) were highly expressed distally (Figure 4D). *In situ* hybridization confirmed proximal *Apccd1* and *Id3* differential expression (Figure 4E). Collectively, these data suggest that the stromal environment is defined by the proximal-distal location along the oesophageal axis. Regional signalling environments are orchestrated by coupling local enrichment of fibroblast subpopulations with a proximal-distal, subpopulation-independent, transcriptional signature.

### Proximal fibroblasts efficiently support oesophageal organoid growth

To investigate the functional implication of regionalized stromal signalling environments, we FACS-isolated oesophageal basal progenitor cells (from all regions) and cultured them with *Pdgfrα*^H2BeGFP^-positive fibroblasts isolated from either the proximal or distal oesophagus (Figure 4F and S3E) ^23^. Both proximal and distal fibroblasts were able to support organoid formation and increasing the number of fibroblasts enhanced oesophageal organoid size independent of the fibroblast origin (Figure 4G). However, organoids cultured with fibroblasts derived from the distal oesophagus were smaller compared to organoids cultured with proximal fibroblasts, indicating that proximal fibroblasts are better at promoting organoid growth (Figure 4H).

Developmental specification of the endoderm requires bidirectional signalling between the epithelium and stroma ^1^. To address if the supportive effect of fibroblasts on organoid growth is dependent on the origin of the epithelial progenitor cells, we co-cultured regionally matched and mismatched oesophageal epithelial basal cells and fibroblasts, respectively (Figure 4I). Adding proximal, but not distal, fibroblasts to proximal epithelium significantly increased organoid size (Figure 4J). Similarly, proximal, but not distal, fibroblasts positively affected the size of distal epithelial organoids (Figure 4K). This suggests that proximal fibroblasts are better than distal fibroblasts at supporting epithelial organoid growth, independent of the epithelial region-of-origin.

### Identification of locally defined oesophageal immune cell repertoires

Our data indicate that fibroblast subpopulations are instrumental in providing defined and regionalized molecular cues that dictate position-specific epithelial basal cell behaviour. However, it is well known that immune cells provide complimentary and interdependent functionality in maintaining tissue homeostasis ^32^. To understand if immune cells contribute to oesophageal regionalization and tissue homeostasis, we subclustered all previously annotated immune cells. We identified a large number of immune cell types, including several subtypes of macrophages, dendritic cells (DCs), T-cells and Langerhans cells. In addition, we identified small populations of Mast cells, Innate lymphoid cells (ILC2s), and natural killer (NK) cells (Figure 5A-B and S4A) (Supplemental Table 10 and 11). All immune cell types localized throughout the oesophageal proximal-distal axis but at varying frequencies (Figure 5C and S4B-D). We found Langerhans, NK-and T-cell populations predominantly in the epithelial fraction, whereas macrophages, dendritic cells and ILC2s were mainly isolated from the oesophageal stroma (Figure 5D and S4D).

**Figure 5.**
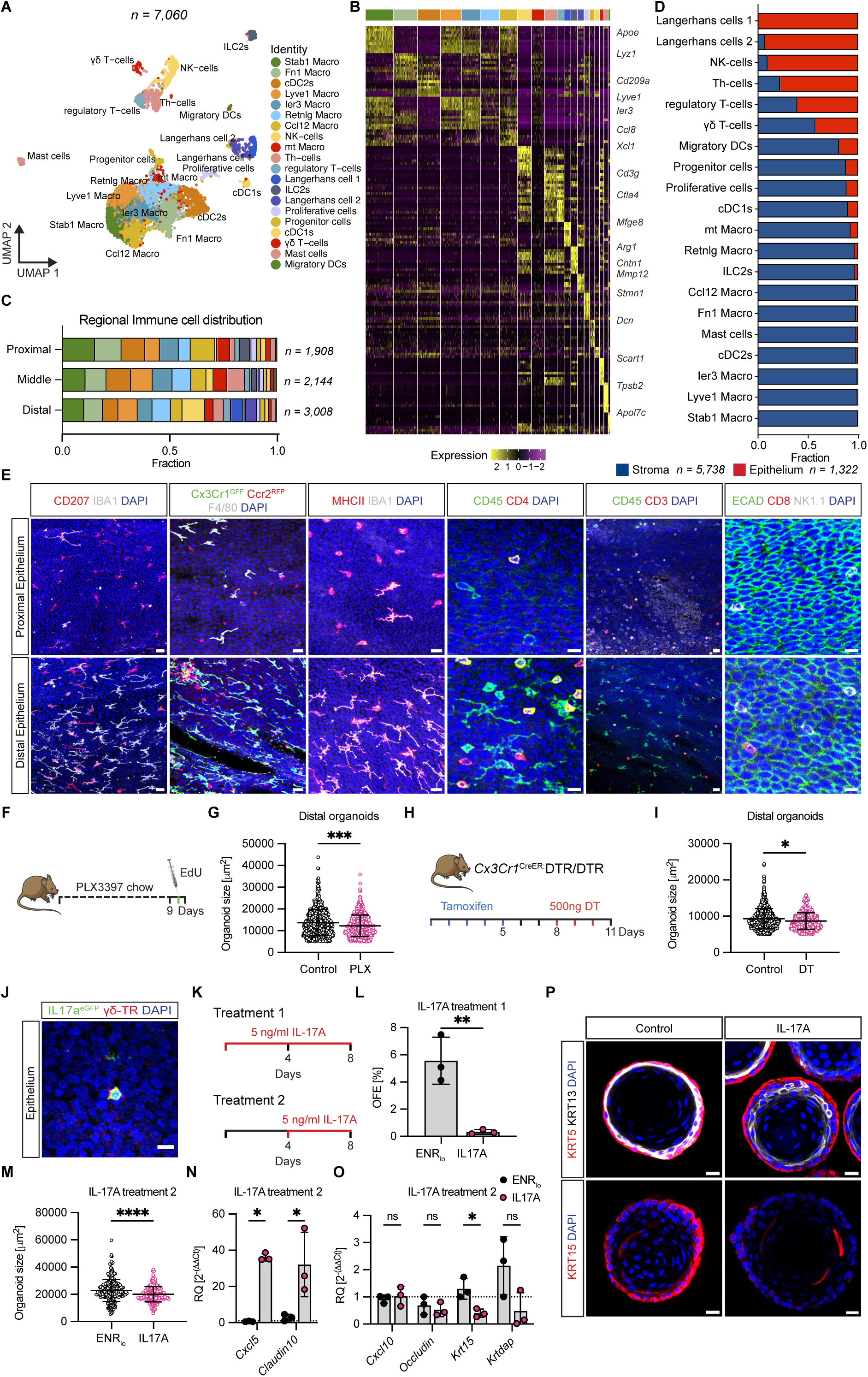
Identification of locally defined oesophageal immune cell repertoires. (A) UMAP of cell identities of subclustered immune cells. (B) Heatmap of top 10 DEGs for all clusters displayed in (A). (C) Distribution of all immune cell populations within each sequenced proximal-distal region. Cell numbers are normalised within respective region, displaying fractions. (D) Quantification the distribution of all immune cell populations across compartment (stroma and epithelium). Fractions are normalised within cell each identity. (E) Immunofluorescent labelling of proximal and distal epithelial whole mounts using CD207 (Langerin), F4/80, Cx3cr1^CreER^:GFP, Ccr2^CreER^:RFP, IBA1, MHCII, CD45, CD4, CD3, CD8, and NK1.1, revealing distal enrichment of intra-epithelial immune cell populations. Scale bar = 20µm and 10µm for the CD45, CD4 staining panel. (F) Experimental setup for *in vivo* depletion of CSF1-dependent immune cells keeping mice on PLX3397 chow *ad libitum* for 9 days. EdU was injected i.p 1 hour before sacrifice. (G) Organoid size derived from the distal epithelium comparing control and PLX3397 chow immune cell depleted mice. n = 3 with 2 mice per n. Each dot represents one organoid. (H) Illustration of experimental setup for *in vivo* depletion of *Cx3cr1* expressing immune cells using Cx3cr1^CreER^:DTR mice. Tamoxifen was injected daily for 5 days (day 1-5) to induce expression of the diphtheria toxin receptor (DTR), after which diphtheria toxin (DT) was administered for 3 days (day 8-10). Mice were sacrificed on day 11. Control mice were treated with tamoxifen, but not DT. (I) Quantification of organoid sizes comparing distal organoids derived from control and DT injected mice. n = 3 with 2 mice per n. Each dot represents one organoid. (J) Immunofluorescent visualization of IL-17A^eGFP^ (green) and ψ8-TR (red) on epithelial whole mounts. Scale bar = 10µm. (K) Schematic illustration of IL-17A treatment. IL-17A was either supplemented when plating cells (day 0, Treatment 1) or from day 4 of culture (Treatment 2). (L) Organoid forming efficiency (OFE) comparing control organoids to organoids grown in the presence of 5ng/mL IL-17A from day 0 onwards. n=3. (M) Quantification of organoid size comparing control to IL-17A when initiated at day 4 (Treatment 2). Each dot represents one organoid. (N) Relative gene expression of *Cxcl5* and *Claudin10* in organoids harvested at day 8, comparing IL-17A Treatment 2 and control. (O) Relative gene expression of *Cxcl10*, *Occludin*, *Krt15* and *Krtdap* in organoids harvested at day 8, comparing IL-17A Treatment 2 and control. (P) Immunofluorescent staining of organoid cross sections, comparing control to IL-17A treated organoids (Treatment 2). Organoids are stained for KRT5 (red, upper panel), KRT13 (white, upper panel), and KRT15 (red, lower panel) and counterstained with DAPI (blue). Scale bar = 20µm. (G, I, M) Two-tailed Kolmogorov-Smirnov test. ns p > 0.05, * p < 0.05, ** p < 0.01, *** p < 0.001. (L) Two-sided ratio paired t-test. ns p > 0.05, * p < 0.05, ** p < 0.01, *** p < 0.001. (N, O) Multiple ratio paired t-test corrected for multiple comparisons with Holm-Šídák method. ns p > 0.05, * p < 0.05, ** p < 0.01, *** p < 0.001.

Using antibodies against specific immune cell populations in epithelial wholemount preparations, we confirmed that the distal epithelium harbours more Langerhans cells (CD207^pos^, MHCII^pos^, IBA1^pos^) than the proximal epithelium (Figure 5E) ^18^. Furthermore, the distinct separation of *Ccr2*^RFP^-positive and *Ccr2*^RFP^-negative *Cx3cr1*^GFP^F4/80-positive intra-epithelial macrophages suggests the turnover of epithelial-resident immune cells ^20^. We also confirmed distal enrichment of intra-epithelial dendritic cells (MHCII^pos^IBA1^neg^) and found that T-cell subpopulations (CD4^pos^, CD3^pos^ and CD8^pos^) as well as NK-cells (NK1.1^pos^) primarily located to the distal oesophageal epithelium (Figure 5E). Using spectral flow cytometry of epithelial and stromal cell fractions we independently confirmed both the identified immune cell types and their regional distributions as indicated by immunofluorescence and single-cell analysis (Figure S4E-F). Taken together, our data suggest that the immune cell landscape is regionalized in two dimensions, coupling a proximal-distal cell gradient with epithelial-stromal cell type-specific separation.

## Immune cells impact oesophageal progenitor cell behaviour

The gradual proximal-distal intra-epithelial increase of Langerhans cells prompted us to probe their functional role in impacting oesophageal progenitor cell behaviour ^18^. To this end, we employed two different *in vivo* ablation strategies. First, we administered PLX3397 (CSF1R antibody-containing chow) for 9 days, depleting CSF1R-dependent immune cells (Figure 5F and S4G) ^33^. Immunofluorescent staining confirmed a reduction in intra-epithelial CD45^pos^/MCHII^pos^ cells after 9 days of PLX3397 treatment while epithelial proliferation was unaltered (S4H). Distal, but not proximal, organoids established from PLX3397-treated, Langerhans cell-depleted, mice were significantly smaller when compared to organoids derived from control proximal and distal oesophagi (Figure 5G and S4I).

Next, we used tamoxifen to induce expression of the diphtheria toxin receptor (DTR) in Cx3Cr1-positive cells (macrophages and intra-epithelial Langerhans cells (Figure S4G)) using the Cx3Cr1-CreER^T2^:DTR mouse line ^34–36^. Subsequent injection of diphtheria toxin resulted in the efficient ablation of CD45^pos^ and DTR-expressing intra-epithelial cells, without significantly affecting epithelial proliferation *in vivo* (Figure 5H and S4J-K). We then FACS-isolated proximal and distal epithelial basal progenitor cells and established organoids. Analogous to PLX3397 administration, genetic ablation of intra-epithelial Langerhans cells reduced the growth of distal, but not proximal organoids when compared to control mice (Figure 5I and S4L), demonstrating that intra-epithelial immune cells impact basal progenitor cell behaviour.

Single-cell sequencing also identified a population of intra-epithelial ψ8-T-cells expressing IL-17A (Figure 5A). We confirmed the presence of rare IL-17A-expressing ψ8-T-cells in the oesophageal epithelium using *IL-17A*^eGFP^ mice (Figure 5J) ^37^. To understand how IL-17A impacts basal progenitor cell properties, we supplemented organoid media with recombinant IL-17A and assessed organoid formation. Addition of IL-17A significantly impaired organoid forming capacity, preventing further mechanistic analysis (Figure 5K-L, Treatment 1). Cell-cell communication prediction using CellChat ^38^ indicated that IL-17A from ψ8-T-cells targets proliferating basal, as well as differentiated suprabasal cells, suggesting that IL-17A impacts organoid formation by impinging on epithelial differentiation (Figure S5M). To probe this possibility, we administered IL-17A at day 4 of organoid culture (Treatment 2), a time point when small, but still expanding, organoids are already formed. We found that organoids growing in the presence of IL-17A were significantly smaller than control organoids (Figure 5K and 5M). Mechanistically, IL-17A induced expression of distinct proinflammatory chemokines (*Cxcl5,* but not *Cxcl10*) and tight junction components (*Claudin 10,* but not *Occludin*), suggesting that IL-17A modulates basal progenitor cell gene expression (Figure 5N-O) ^39^. In addition, we found that IL-17A treatment significantly reduced the basal progenitor marker *Krt15* as well as supressed *Krtdap* (a marker for differentiation). Immunofluorescent analysis of organoids confirmed that suprabasal KRT13 as well as basal KRT15 were reduced after IL-17A exposure (Figure 5P). Taken together, these experiments suggest that ψ8-T-cells impact oesophageal epithelial barrier homeostasis ^40,41^.

## Deconvoluting the cellular architecture mediating regional signalling activity

Regionalized cellular architecture likely corresponds to local molecular and signalling activities. To deconvolute the proximal-distal signalling landscape, we revisited CellChat. Since fibroblasts are well-established mediators of epithelial progenitor behaviours, we focused our efforts on understanding epithelial-fibroblast communication. We predicted the number of interactions between three main epithelial cell groups (Proliferating, Basal and Suprabasal) and fibroblasts and found that communication between all populations was increased in the distal compared to the proximal oesophagus, suggesting a distal enrichment of cell-to-cell interaction potential (Figure S5A-B). Including all epithelial and fibroblast subpopulations confirmed that the inferred interaction strength was elevated in the distal compared to the proximal oesophagus (Figure S5C). Taken together, these data indicate a complex and regionalized oesophageal signalling landscape.

## Exploring proximal-distal signalling gradients

Focusing on identifying signalling pathways differentially engaged along the oesophageal axis, we compared the inferred proximal and distal epithelial-to-fibroblast communication landscape. We identified a non-uniform, predicted activity of multiple signalling pathway (Figure S5D), where Neuregulin (NRG) signalling was enriched in the proximal and IGF and BMP signalling in the distal oesophagus. Interestingly, WNT signalling was predicted to be active in both proximal and distal oesophagi (Figure 6A). These data suggest a mechanism whereby differential engagement of signalling pathways along the oesophageal axis allows for fine-tuning of basal progenitor cell states and behaviour.

**Figure 6.**
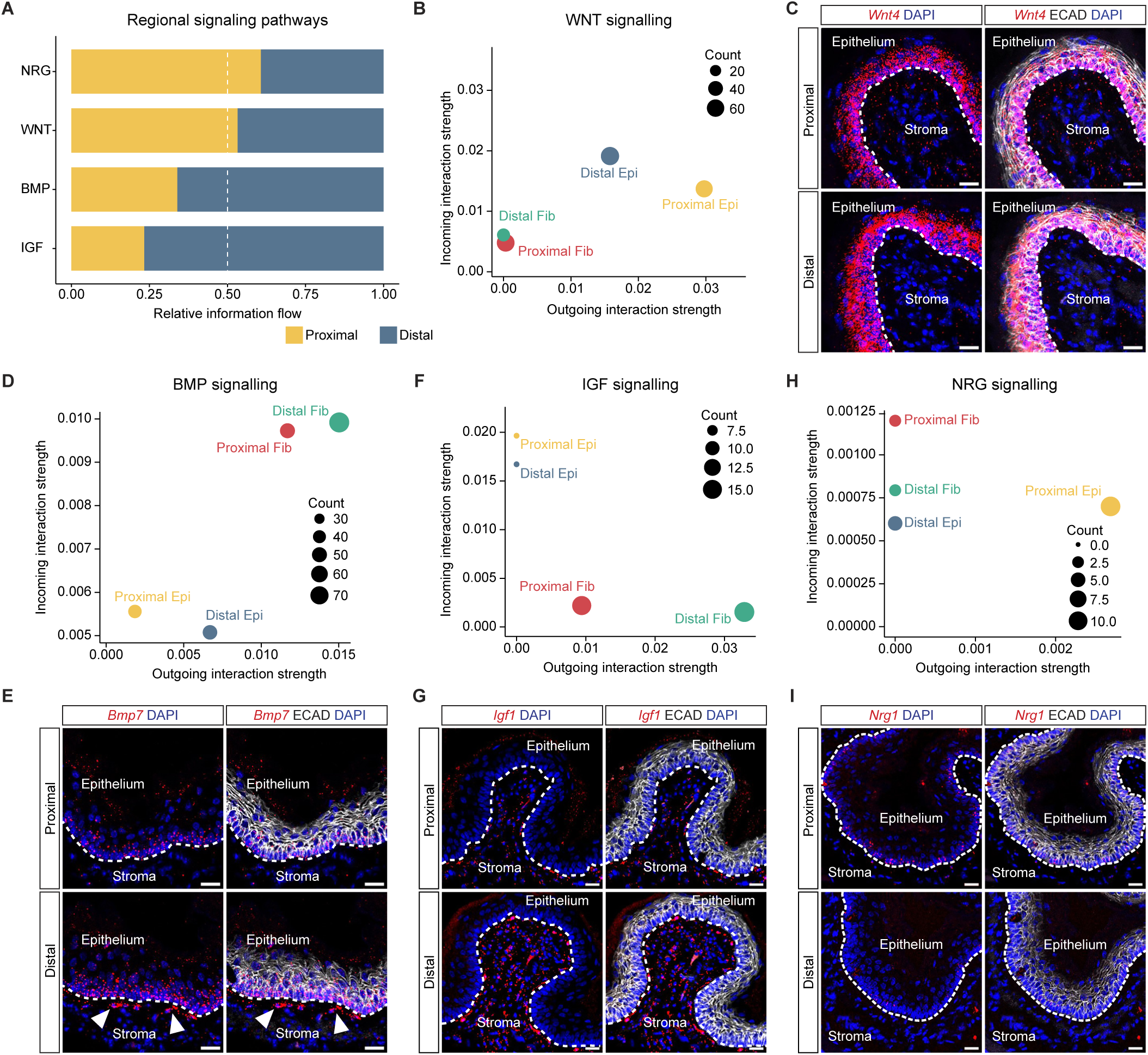
Exploring proximal-distal signalling gradients. (A) Information flow (sum of communication probabilities among pairs of cell groups) of selected signalling pathways comparing datasets encompassing all proximal and distal cells within the single-cell dataset. (B, D, F, H) Comparison of the outgoing and incoming interactions strengths between proximal-distal fibroblasts and proximal-distal epithelial cells, focusing on (B) WNT-, (D) BMP-, (F) IGF- and (H) NRG-signalling. (C) *In situ* hybridization for *Wnt4* (red), enriched in the epithelium. (E) *In situ* hybridization for *Bmp7* (red). White arrows mark *Bmp7*-positive stromal fibroblasts in close contact with the epithelium. (G) *In situ* hybridization for *Igf1* (red), enriched in the distal, compared to the proximal, stroma. (I) *In situ* hybridization for *Nrg1* (red), expressed in the proximal, but not distal, epithelium. All *in situ* hybridizations (C, E, G, and I) are performed on cross-sections of the proximal and distal oesophagus and counterstained with E-CADHERIN (ECAD, white) and DAPI (blue). Dotted lines mark the epithelial-stromal border. Scale bar = 20µm.

Although WNT signalling is implicated in renewal and tumour formation of several epithelia ^42–44^, the function of WNT in the oesophagus is not clear ^45,46^. Closer analysis of the predicted WNT signalling activity revealed that proximal as well as distal epithelial cells appeared to be both the agent and recipient of WNT signalling, suggesting a largely intra-epithelial signalling mechanism (Figure 6B, S5E-F). Exploring the single-cell data, we found high *Wnt4* and lower *Wnt10a* and *Wnt5a* expression in the oesophageal epithelium (Figure S5G). *In situ* hybridization confirmed high epithelial expression of *Wnt4* (canonical and/or non-canonical) ^47,48^, suggesting that epithelial WNT signalling is predominantly WNT4-dependent (Figure 6C). When assessing the canonical WNT-target gene *Axin2,* we found overall low epithelial expression, although *Axin2* levels were elevated in the distal compared to the proximal epithelium (Figure S5H). Interestingly, the non-canonical WNT ligand *Wnt5a* displayed a similar proximal-distal epithelial expression pattern and both *Axin2* and *Wnt5a* were enriched in the stroma (Figure S5H), suggesting that the function of WNT signalling could extend beyond autocrine epithelial signalling (Figure 6B).

BMP signalling is a known mediator of oesophageal progenitor cell differentiation ^49,50^. We found that epithelial-directed BMP signalling predominantly originates from distal fibroblasts (Figure 6D and S5E-F). To identify a stromal BMP-signalling source, we analysed all fibroblast subpopulations and verified distally enriched *Sfrp2*-fibroblasts as a predicted signalling hub (Figure S5I), expressing several BMP ligands (*Bmp2*, *Bmp4*, *Bmp5* and *Bmp7*). *In situ* hybridization confirmed *Bmp7* expression in distal stromal cells adjacent to the oesophageal epithelium (Figure 6E).

We then focused on IGF signalling since this pathway not only promotes growth of epithelial cells but is also linked to oesophageal disease ^51–53^. We found that IGF signalling originated from distal fibroblasts and signalled towards the epithelium (Figure 6F and S5E-F). Although *Igf1* was expressed by several fibroblast subpopulations, distal fibroblasts generally exhibited elevated *Igf1* expression compared to proximal fibroblasts (Figure S5J). Interestingly, BMP-producing *Sfrp2*-fibroblasts displayed the highest expression of *Igf1.* The IGF1 receptor (*Igf1r*) was expressed in all epithelial basal subpopulations irrespective of location, indicating that stromal IGF1 could target basal oesophageal progenitor cells (Figure 6I and S5K). *In situ* localization of *Igf1* mRNA confirmed a distal stromal enrichment (Figure 6G).

NRG1 belongs to the EGF growth factor family and mediates intestinal stem cell proliferation and regeneration potential ^54,55^. We explored oesophageal NRG1-signaling, predicted to originate in the proximal oesophageal epithelium (Figure 6H). We assessed *Nrg1* and cognate *Erbb*-receptor expression and observed *Nrg1* to be enriched in proximal basal progenitor cells (Basal 1 and Basal 2), whereas the *Erbb2* and *Erbb3* receptors were expressed by both basal and suprabasal epithelial cells irrespectively of their proximal-distal location (Figure S5L). In addition, *Nrg1* mRNA localized to the proximal, but not distal, oesophageal epithelium (Figure 6I). Altogether, our work reveals how regional differences in cellular and molecular architecture result in local signalling environments, likely reflecting local progenitor cell states and mechanisms of tissue homeostasis.

### Local cellular architectures shape the oesophageal signalling landscape

Cell-cell communication analysis suggested that the proximal and distal oesophageal signalling landscapes are distinct, but did not to provide insights into either the functional role of specific signalling pathways or if basal progenitor cells display a regionalized response to specific signalling ligands. To this end, we isolated proximal and distal basal progenitor cells and evaluated organoid forming efficiency and organoid size after six days of culture in the presence of signalling modulators. Organoid forming efficiency of both proximal and distal progenitor cells was significantly decreased in the presence of distally enriched BMP4 (Figure 7A). Enhancing endogenous BMP activity by reducing the concentration of NOGGIN recapitulated BMP4 treatment (Figure S6A-B), supporting previous work demonstrating that BMP inhibition is required for organoid formation, acting by preventing differentiation ^49,50^. In contrast to BMP4, IGF1 enhanced proximal and distal organoid size, but not organoid forming efficiency (Figure 7B and S6B) supporting an epithelial growth-promoting role of IGF1 normally enriched in distal fibroblasts. NRG1, expressed by proximal basal epithelial cells enhanced the size of both proximal and distal organoids, whereas only proximal, and not distal, organoid forming efficiency was positively affected by the addition of NRG1 (Figure 7C-D). Lastly, we evaluated regional effects of modulating WNT signalling *in vitro*. Although preventing epithelial WNT-secretion in either proximal or distal basal cells using IWP2 (a PORCN inhibitor) ^56^ did not significantly alter neither organoid formation nor growth (Figure S6D-E), addition of WNT5A reduced organoid forming efficiency, but not size, of both proximal and distal organoids (Figure 7E and S6F). These data indicate a role for non-canonical, rather than canonical, WNT signalling in restricting basal progenitor cell behaviour. We demonstrate that although both proximal and distal basal cells have equal potential to respond to signalling cues, with the notable exception of NRG1-sensitivity, the *in vivo* regional cellular architecture efficiently restricts signalling to specific oesophageal epithelial regions (Figure 7F).

**Figure 7.**
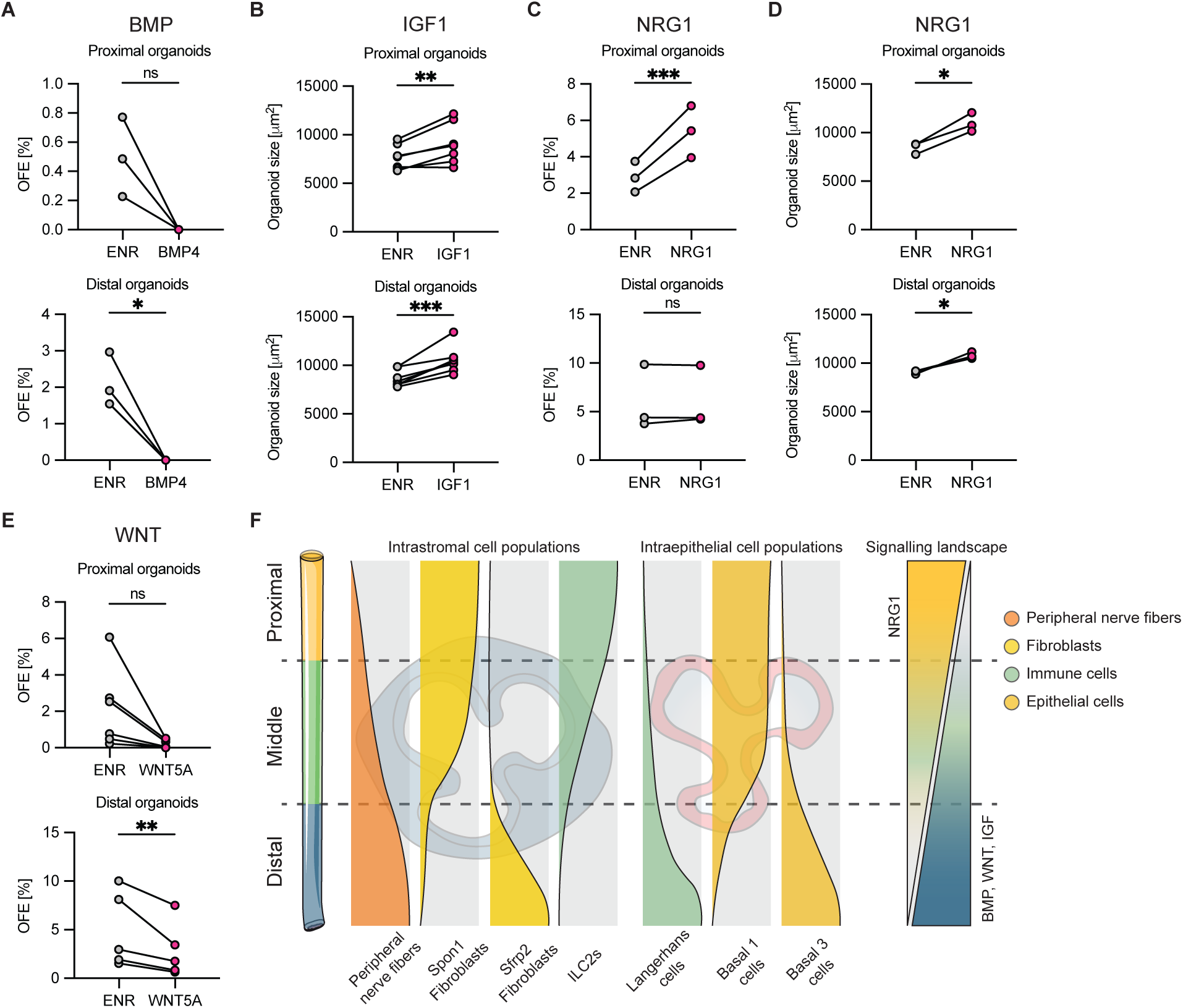
Local cellular architectures shape the oesophageal signalling landscape. (A) Quantification of proximal and distal organoid forming efficiency (OFE) comparing control medium and medium containing 100ng/mL BMP4. (B) Quantification of organoid size in control medium and medium supplemented with 250ng/mL IGF1. (C-D) Quantification of organoid forming efficiency (C) and size (D) in control medium and medium supplemented with 100ng/mL recombinant NRG1. (E) Quantification of organoid forming efficiency in the presence of 250ng/mL WNT5A. All quantifications were performed at day 6 of culture. For size quantifications each dot presents average size of all organoids. n=3-5. (F) Graphical summary illustrating the regionalized oesophageal stromal and epithelial cell distributions as well as signalling gradients. (A-E) Two-sided ratio paired t-test. ns p > 0.05, * p < 0.05, ** p < 0.01, *** p < 0.001.

## Discussion

The oesophagus is an essential beginning of the gastrointestinal tract. Despite that mortality due to oesophageal cancer is significantly higher than for other gastrointestinal cancers, little is known about the cellular and molecular mechanisms that drive oesophageal disease. Clinical observations suggest that oesophageal cancers, as well as inflammatory disease, tend to spare the proximal oesophageal tube, while preferentially targeting middle and distal regions ^14–17^, likely reflecting how different oesophageal regions respond to external challenges. Herein, we provide a complete, regionally resolved, cell atlas spanning the entire oesophageal axis and encompassing all major cell types. We combine in-depth transcriptomic characterization with *in vivo* proof-of-concept models to demonstrate functional cellular as well as molecular oesophageal regionalization. This work provides a framework for future research focused on understanding local disease occurrence in the oesophagus.

The oesophagus is traditionally perceived as a largely uniform tube. We report that the tissue architectures of the proximal and distal oesophagus are macroscopically distinct and reflect the prevalence of well-established cell types known to impact epithelial progenitor cell states and tissue homeostasis (Figure 1). Regionalized Smart-seq3 single-cell sequencing revealed non-uniform axial distribution of subpopulations of epithelial cells and fibroblasts and demonstrated the existence of proximal-distal differences in immune cell types and frequencies. We identified five basal progenitor cell subpopulations and unveiled a gene signature expressed in stretches of basal progenitor cells in an alternating circumferential pattern in the distal basal epithelium (Figure 3). Although it is not clear what the uneven circumferential distribution of this basal subpopulation represents, it is tempting to speculate that food-induced expansion of the folded oesophageal tube requires a subset of progenitor cells that can efficiently sustain or respond to changes in mechanical forces.

Developmental patterning of the intestinal tract establishes gradients of signalling effectors maintained in the adult. To understand this aspect of oesophageal regionalization, we compared all proximal to distal epithelial cells and identified a transcriptional signature representing the proximal-distal position along the oesophageal tube (Figure 2). Thus, transcriptional signatures related to proximal-distal cell location thus impact all epithelial cell states. However, regional differences in both organoid forming efficiency as well as gene expression profiles are lost upon passaging of organoids. This indicates that although distinct distal epithelial cell states exist, the epithelium is defined by transcriptional programs likely established by local epithelial-stromal communication.

Focusing on non-epithelial cells to understand regionalized epithelial progenitor behaviour, we identified eight subpopulations of fibroblasts unevenly distributed along the oesophageal axis (Figure 4). We demonstrated how *Sfrp2*-positive fibroblasts enable distal BMP- and IGF-signalling activities, representing a functional equivalent to the BMP^high^ fibroblasts described in the intestine ^12,13^. In addition, we determined a subpopulation-independent, transcriptional stromal profile where signalling molecules linked to WNT, BMP/TGFβ and HH signalling were associated with either proximal or distal fibroblasts. Organoid-fibroblast co-cultures demonstrated that while both proximal and distal fibroblasts were permissive to organoid formation, organoids kept in the presence of proximal fibroblasts were significantly larger than organoids cultured with distal fibroblasts. Importantly, proximal fibroblasts promoted growth of proximal and distal organoids alike, reiterating the importance of the stroma in establishing epithelial cell behaviour.

Finally, we identified both stromal and intra-epithelial immune cell types (Figure 5). We found proximal-distal gradients of intra-epithelial Langerhans, NK- and T-cells, mirroring findings from the human oesophagus ^18^ and indicating that the immunological response to inflammatory agents challenging the epithelial barrier is regionalized. *In vivo* depletion of macrophages and distally enrichment of intra-epithelial Langerhans cells decreased growth of organoids derived from distal, but not proximal progenitor cells, unveiling a region specific immune-epithelial cell circuit. In addition, we identified a scarce population of intra-epithelial IL-17A-positive ψ8-T-cells strategically located to impact oesophageal tissue response to inflammatory challenges.

To understand if the regionalized cellular architecture corresponds to differential engagement of signalling pathways along the oesophageal axis, we inferred cell-cell communication networks along the oesophageal axis. We found that proximal-distal and distal-proximal oesophageal signalling gradients are the norm rather than the exception (Figure 6 and 7). Focusing on key pathways, we discovered that BMP- and IGF-signalling emanated from the stroma, whereas *Nrg1* is specifically localized to the proximal epithelium. We demonstrated that NRG1 can signal in an autocrine manner, affecting organoid forming efficiency of proximal, but not distal, basal progenitor cells. Finally, we observed *Wnt4* to be highly expressed in the oesophageal epithelium, although signs of epithelial canonical WNT signalling were low, suggesting that oesophageal tissue homeostasis was largely maintained by non-canonical WNT signalling. How gradients of signalling pathways crosstalk, synergize or antagonize remains to be determined, but is likely to contribute additional complexity to oesophageal tissue maintenance.

We report that oesophageal epithelial homeostasis is regionalized, providing mechanistic understanding for future directed interventions targeting oesophageal disease.

### Limitations of study

This work explores the mouse oesophageal axis. Cell states and functions are deduced from transcriptional profiles and cell-cell communication predictions in combination with *in vivo* genetic lineage tracing/depletion models and organoid co-cultures. Due to limited resources, we were not able to experimentally probe the *in vivo* function of each identified subpopulation and cell type. This study does not consider how extrinsic factors such as mechanical forces, pH and microbiome impact gene expression.

## Methods

### Animal husbandry and animal procedures

Animals were bred and housed in pathogen-free conditions according to the recommendations of the Federation of European Laboratory Animal Science Association. All animal experiments were approved by the appropriate ethical review board (Stockholms norra djurförsöksetiska nämnd) (ethical permits 14051-2019 and 735-2021). All experiments were initiated when mice were between 10 - 14 weeks old. C57Bl/6 mice were purchased from Charles River (C57BL/6J) or bred locally. Sox2-CreER^T2^ and PdgfRa-H2BeGFP lines are available at JAX (Strain #017593 and #007669 respectively) ^28,57^. Cx3Cr1^GFP^ ^58^; Ccr2^RFP^ ^59^ and Cx3cr1^CreER-eYFP^ ^35,36^ were bred locally. IL-17A^IRES-eGFP^ ^37^ knock-in tissue was kindly provided by E. Villablanca at Karolinska Institutet. Except ear clipping for genotyping purposes, no previous procedures were performed on experimental animals.

#### Genotyping

Ear biopsies were lysed overnight at 55°C in DirectPCR lysis reagent (BioSite, #250-102-T) supplemented with 0.1 mg/ml Proteinase K (Thermo Fisher Scientific, #EO0492). Lysis was stopped by heat inactivation at 85°C for 45 min. Amplification was conducted in 1x PCR buffer supplemented with 4 mM MgCl2, 1 mM dNTPs, and 0.5 U/µl Taq Polymerase (Invitrogen, #10342046). Additionally, target specific primers were added at 0.4 µM (IDT). PCR products were analysed on a 1-2 % agarose gel (VWR, #732-2789P) prepared in 1X TAE buffer (VWR, #K915) containing 1X GelRed solution (VWR, #41003).

#### Tamoxifen and EdU administration

Ccr2^RFP^Cx3cr1^eGFP^ mice were recombined using i.p injections of 200µl of 1% Tamoxifen (W/V, Sigma-Aldrich, #T5648) in corn oil (Sigma-Aldrich, #C8267) 4 times within 5 days. Cx3cr1^CreER^:DTR/DTR mice were recombined using i.p injections of 100µl of 2% Tamoxifen (W/V, Sigma-Aldrich, #T5648) in corn oil for 5 consecutive days. PDGFRα^H2BeGFP^ mice were recombined using i.p injections of 100µl of 2% Tamoxifen in corn oil for 3 consecutive days. To assess proliferation, 100μl of 1 mg/ml EdU (Sigma-Aldrich, #900584) in PBS was injected i.p between 6 and 7 am for 1 hour.

#### Macrophage depletion using PLX3397 and diphtheria toxin

The CSF1R- and c-Kit antibody Pexidartinib (PLX3397) was formulated into A04 standard diet (Safe Nutrition Service, France) at 290ppm and was administered *ad libitum* for 9 consecutive days. Identical diet without PLX3397 was used as control. To achieve depletion of *Cx3cr1* expressing cells 100µl of diphtheria toxin (500ng/mL) was injected i.p for 3 consecutive days 2 days after the last Tamoxifen injection.

### Immunostaining

Oesophagi were dissected and cut in regional segments. Subsequently, the segments were either snap frozen in Tissue-Tek® O.C.T. Compound (Sakura, #4583) or fixed with 4% formaldehyde solution (V/V, Sigma, #252549), kept overnight in 20% sucrose in PBS (W/V, Sigma-Aldrich, #84100), finally embedded in O.C.T and kept in -80°C until sectioned on glass slides. Sections were cut at 10µM thickness with a cryotome (ThermoFisher). For immunostainings, sections were blocked in blocking buffer containing donkey serum (2.5% V/V, Jackson Immuno, #017-000-121), goat serum (2.5% V/V, JacksonImmuno, #005-000-121), BSA (1% W/V, Sigma, #A4503), and Triton X-100 (0.3% V/V, Merck, #93443) in PBS for 1 hour. Primary antibodies were incubated overnight in blocking buffer. After 3 times washing in PBS, secondary antibodies (1:1000, Jackson Immuno) were incubated for 1 hour at RT in blocking buffer. Following 3 times washing with PBS, sections were incubated with DAPI (0.25 µg/mL, Thermo Fisher Scientific, #D1306) in PBS. Finally, sections were mounted with ProLong™ Gold Antifade Mountant (Invitrogen, #P36930). Images were acquired with either a Zeiss AxioImagerZ2 or Zeiss LSM980 microscope.

#### Determining αSMA muscle layer thickness

At least three cross-sections from five individual mice were quantified for the proximal or distal oesophagus, respectively. All specimens were subjected to the same staining procedure and image acquisition was performed with the same microscope settings. At ten to twenty randomly chosen locations a line perpendicular to the oesophageal epithelium was manually drawn spanning the entire thickness of the αSMA-positive layer. Subsequently, the ImageJ (v2.9.0/1.53t) *Measure* function was used to determine the length of the lines in µm and the average of all measurements was calculated per section.

#### Fluorescent intensity quantification

LRIG1 intensity was quantified using ImageJ2 (v2.90/1.53t) ^60^. Area integrated intensity and mean grey value of at least six manually selected regions from three images were quantified from proximal and distal cross-sections of the oesophagus, respectively. Additionally, three areas without fluorescent signal were measured to calculate background signal intensity. Corrected total cell fluorescence (CTCF) was then calculated using the formula:

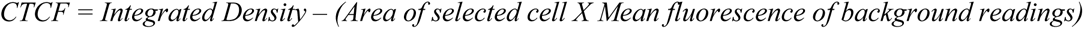

#### KLF4-positive cell quantification

At least three cross-sections of the proximal and distal oesophagus were analysed from three mice, respectively. All specimens were subjected to the same staining procedure and image acquisition was performed with the same microscope settings. ImageJ was used for quantifications after the unspecific background signal was removed, and the same arbitrary threshold was applied to all images. KLF4-positive and KLF4-negative cells were manually counted.

### Whole mount immunostaining

Whole mount immunostainings were conducted as described previously ^61^. In brief, oesophagi were dissected and the muscularis externa mechanically removed using forceps. Subsequently, the oesophagus was longitudinally opened and cut into approximately 5 mm segments. Specimen were incubated in 5mM EDTA (VWR, #E522) at 37°C for 2 - 3 hours to be able to separate the epithelium and stroma. Tissue pieces were fixed for 30min in 4% Formaldehyde (V/V, Sigma-Aldrich, #252549) in PBS at 4°C. Tissue pieces were submerged in whole mount blocking buffer (2.5% donkey serum, 2.5% goat serum, 1% BSA, 0.5% Triton X-100 in PBS) for 1 hour at RT. Blocked tissue pieces were incubated in primary antibodies diluted in whole mount blocking buffer overnight or for up to three days at 4°C. Following, tissue pieces were washed in 0.2% Tween 20 in PBS (V/V, Bio-Rad, #1706531) for 2 - 3 hours exchanging washing buffer approximately every 30min. Corresponding secondary antibodies (Jackson Immuno, 1:1000) were incubated for 3 hours at RT in blocking buffer or overnight at 4°C. All secondary antibodies were anti-Goat or anti-Donkey Alexa Fluor-488/Cy3/647). Finally, tissue segments were incubated overnight at 4°C in DAPI (0.25µg/mL, Thermo Fisher Scientific, #D1306) in PBS before spread and carefully flattened on object slides and mounted in ProLong™ Gold Antifade Mountant (Invitrogen, #P36930). Images were acquired with either a Zeiss LSM980 or Leica Stellaris 5.

The following antibodies were used for immunostainings at given dilutions:

Rat anti-CD3 (BioLegend, clone 17A2, #100201, 1:100), Rat anti-CD4 (BioLegend, clone GK1.5, #100401, 1:100), Rat anti-CD8 (Bio-Rad, #MCA1768, 1:100), Armenian-Hamster anti-ψ8T-cell receptor (BD Biosciences, clone GL3, #553175, 1:50), Rat anti-MHCII (BD Pharmingen, clone M5/114.15.2, #556999, 1:100), Rat anti-Langerin (CD207) (eBioscience, clone eBioL31, #14-2075-82, 1:100), Rabbit anti-IBA1 (Wako Chemicals, #019-19741, 1:500), Rat anti-F4/80 (Bio-Rad, clone CI:A3-1, #MCA497GA, 1:50), Rabbit anti-CD45 (Abcam, #ab10558, 1:500), Rat anti-CD45 (Invitrogen, clone 30-F11, #14-0451-82, 1:100), Mouse anti-αSMA (Sigma-Aldrich, clone 1A4, #A2547, 1:1000), Rat anti-CD31 (BD Pharmingen, clone MEC13.3, #550274, 1:100), Mouse anti-CD31 (Thermo Fisher Scientific, clone 2H8, #MA3015, 1:1000), and Rabbit anti-beta III tubulin (TUJ1) (Abcam, #ab18207, 1:500).

### Imaging analysis

Either an inverted Zeiss LSM980 laser-scanning confocal microscope equipped with four lasers (405nm, 488nm, 561nm, 640nm) or an inverted Leica Stellaris 5 with a white laser configuration were used for image acquisition. Typical settings used a 20X oil objective, optimal pinhole, a scan speed of 600 Hz, a line average of four, bi-directional scanning, step sizes of 2-4 μm, and a resolution of 1024 x 1024 pixels. Images were reconstructed based on optical sections using ImageJ2 (v2.90/1.53t) or Imaris (v9.0.1, Oxford Instruments). Quantifications were conducted in Imaris using semi-automated spot detection assuming a 6 μm radius of DAPI and EdU positive nuclei and an arbitrary quality threshold kept constant within experiments was applied. The threshold was chosen on visual inspection to be above the background level for the DAPI and EdU staining, respectively.

#### Sox2^CreER^:YFP-positive clone size quantification

Three months after Tamoxifen recombination basal cell clone sizes were manually counted based on whole mount images of proximal and distal oesophageal epithelium. CD49f was used as a counterstaining to ensure localisation to the basal layer and distinguish single cells. 764 clones from the proximal and 783 clones from the distal oesophagus were counted from three mice (176, 339, and 249 clones from proximal; 286, 349, and 148 clones from distal).

#### Quantification of TUJ1- and CD31-positive areas

Images were reconstructed based on optical sections using ImageJ2 (v2.90/1.53t) and maximum intensity projections of the acquired z-stacks performed. Oesophageal tissue was selected using the freehand selection tool and the positive area calculated based on an arbitrary threshold applied to all images.

### qPCR

Ice-cold PBS (Gibco, #14190250) was used to isolate organoids from Matrigel (Fisher Scientific #356231) domes. Subsequently, organoids were washed 3 times with ice-cold PBS (Gibco, #14190250) and kept on ice for the entire isolation procedure. RNA was isolated using the RNeasy Mini Kit (QIAGEN, #74106) following manufacturer’s instructions. Briefly, organoids were resuspended in 350µl RLT buffer and vortexed for 15-20s to disrupt the organoids. 350µl 70% (V/V) ethanol were added and the sample spun at > 8000 x g for 15 s. Subsequently, columns were washed once with 700ul RW1 buffer. Afterwards, samples were washed twice with 500µl RPE buffer. Finally, RNA was eluted in 40µl DNase/RNase free water. After quality check using Nanodrop and including samples with a 260/30 ratio >1.8, 500ng total RNA was used for cDNA synthesis using SuperScript™ VILO™ cDNA Synthesis Kit (Invitrogen, #11754250) following manufacturer’s instructions. Diluted cDNA was used together with selected primer pairs and Power SYBR® Green PCR Master Mix (Thermo Fisher Scientific, #4368702) on a ViiA7 device (Applied biosystems) using a custom standard 2-step protocol. *Hprt* and *18srRNA* were used as internal controls.

### RNAscope *in situ* hybridization

RNAscope was performed according to the manufacturer’s protocol for the RNAscope® Multiplex Fluorescent Detection Kit v2 (ACD, 323110). In brief, fixed frozen sections were allowed to equilibrate to RT and washed in 1X PBS for 5min. The slides were transferred to 1X target retrieval solution (ACD, 322000) at 100°C and kept in for 5min. Afterwards, slides were washed in MilliQ and 100% ethanol for 3min, respectively. The slides were treated with hydrogen peroxide (ACD, 322335) for 10min. Slides were washed twice for 1min in MilliQ. Subsequently, slides were treated with Protease III (ACD, 322337) for 30min at 40°C in the in the HybEZ™ II Oven (ACD, # 321720). Slides were washed in MilliQ and 1X RNAscope probes were applied. When applicable RNAscope probes were diluted 50 times in a -C1 probe. Otherwise, RNAscope probes were diluted 50 times in RNAscope dilution buffer. RNAscope probes against Axin2 (ACD, #400331-C2), Wnt4 (ACD, #401101-C2), Bmp7 (ACD, #407901), Igfbp2 (ACD, #405951), Igfbp5 (ACD, #425731-C2), Smoc2 (ACD, #318541-C3), Wnt5a (ACD, #316791-C3), Apcdd1 (ACD, #425701), Id3 (ACD, #445881-C2), Pappa (ACD, #443921) and custom made Tmprss11a (ACD, #1232761-C1) were applied to the slides and incubated for 2 hours at 40°C in the HybEZ Oven. The slides were washed using 1X wash buffer (ACD, #310091) and the amplification reagents AMP1, AMP2 and AMP3 were applied for 30, 30, and 15 minutes, respectively at 40°C with two washes of 5min in RNAscope wash buffer after each incubation. If applicable, slides were incubated with HRP-C1 for 30min at 40°C in the HybEZ Oven. After two washes of 5min in 1X RNAscope wash buffer slides were incubated with 1:1500 Cyanine 3 amplification reagent (PerkinElmer, #FP1170012UG), or 1:1000 Cyanine 5 amplification reagent (PerkinElmer, #FP117024UG) in TSA buffer (ACD, #322809) for 30min at 40°C in the HybEZ Oven. Upon two more 5min washes in 1X RNAscope wash buffer HRP blocker solution was applied and incubated for 15min at 40°C in the HybEZ Oven. If applicable, the same steps were repeated for the HRP-C2 or HRP-C3 solutions with the respective complementing Cyanine amplification reagent. Finally, a final 5min MilliQ wash was performed. In case of additional immunostainings, slides were blocked with blocking buffer (1% BSA, 2.5% NDS, 2.5% NGS, 0.3% Triton X-100 in PBS) for 30min at RT. Subsequently, slides were incubated with primary antibody overnight. Following, 3 washes with 1X PBS, secondary antibodies were applied for 1 hour at RT. All secondary antibodies used were anti-Goat or anti-Donkey Alexa Fluor-488/Cy3/647 (Jackson Immuno). Subsequently, slides were counter-stained with DAPI (0.25µg/ml, Thermo Fisher Scientific, #D1306) for 5min at RT. Finally, after a 5min wash in 1X PBS slides were mounted on object slides using ProLong™ Gold Antifade Mounting media (Invitrogen, #P36930). Images were acquired using a Zeiss LSM980 laser-scanning confocal microscope equipped with four lasers (405nm, 488nm, 561nm, 640nm).

### Oesophageal cell isolation

Mice were sacrificed using CO_2_. For single-cell sequencing experiments mice were perfused with at least 50ml ice-cold DPBS (Gibco, #14190250). The oesophagus was dissected and the muscularis externa mechanically removed using forceps under a dissection microscope. The stripped oesophagus was longitudinally cut open and incubated in 0.5mg/ml Thermolysin (Sigma-Aldrich, #T7902) at 37°C for 15min. Subsequently, the stroma and epithelial layer were carefully separated under a dissection microscope and cut into 3-4 equally sized tissue pieces. Afterwards, the tissue was thoroughly minced and incubated in 0.25mg/ml Liberase^TM^ (Sigma-Aldrich, #5401127001) and 0.25mg/mL DNase I (Sigma-Aldrich, #11284932001) in HBSS (Gibco, #14175-129) at 37°C for 30min or 60min for stroma and epithelial layer, respectively, pipetting thoroughly up and down every 15min using a 200µl pipette tip. Afterwards, cells were washed twice in 1% FBS in PBS (V/V, Sigma-Aldrich, #7524) and kept on ice.

#### Oesophageal cell isolation for single-cell RNA sequencing

For single-cell sequencing experiments samples were additionally run on a gentleMACS™ Dissociator (Miltenyi Biotec, #130-093-235) with the custom program h_skin_01 (Duration: 36 s, rpr: 116). Three independent runs were performed. The first experiment is special in performing cell dissociation using incubation with Collagenase (125mg/ml, 1 hour, 37°C, pipetting every 15min, Sigma-Aldrich, #C2674) and subsequently Trypsin without EDTA (2.5% in HBSS, 12min, 37°C, Thermo Fisher Scientific, #15090046) after separation and mincing of the stroma and epithelial layer. Moreover, male and female mice were kept separately in the first experiment to identify potential sex specific cell populations. However, no differences were observed. For sequencing replicates two and three the previously described procedure for cell isolation was conducted.

### Cell sorting

Cells were sorted using a FACS AriaIII (BD) equipped with 4 lasers (405nm, 488nm, 532nm, 640nm) and a 100µm nozzle. Cells from the stroma were sorted gating on single GFP-positive cells from Pdgfrα^H2Be-GFP^ mice or using anti-CD45-PE/Cy7 (BD Pharmingen, clone 30-F11, #552848) antibodies. Cells from the epithelial layer were sorted gating on single positive for CD326-PE/Cy7 (BioLegend, clone DECMA-1, #147310) and CD104-AF647 (BioLegend, clone 346-11A, #123608) while simultaneously excluding CD45-FITC (BioLegend, clone 30-F11, #103107) positive cells. In both cases, dead cells were removed during acquisition using SYTOX™ Blue Dead Cell stain incubated for at least 3min on ice prior to acquisition (Invitrogen, #S34857, 1:10.000).

#### Cell sorting for single-cell RNA sequencing

For single-cell RNA sequencing experiments cells were washed and stained with anti-CD45-PE/Cy7 (BD Pharmingen, clone 30-F11, #552848) and SYTOX™ Green Ready Flow™ Reagent (Invitrogen, #R37168, 1:10.000) before sorting into 384-well plates containing lysis buffer (described below) using a BD FACSMelody^TM^ (BD Biosciences) instrument with a 100µm nozzle. A gating strategy to specifically sort for CD45-positive and CD45-negative cells was further performed on all experiments.

### Spectral flow cytometry

After cell isolation according to the procedure above, cells were washed twice with 1% FBS (V/V, Sigma-Aldrich, #7524) in PBS and the remaining cell pellet was resuspended in PBS containing the following antibodies and stainings:

Violet - Live/Dead (Thermo, #L23105), CCR2-BV421 (clone 475301, BD Pharmingen, #747963), Ly6C-BV510 (clone HK1.4, BioLegend, #128033), TCR-gd-BV605 (clone GL3, BioLegend, #118129), NK1.1-BV711 (clone PK136, BioLegend, #108745), CD11c-BV785 (clone N418, BioLegend, #117336), MHCII-488 (clone M5/114.15.2, BioLegend, #107615), CD207-PE (clone 4C7, BioLegend, #144203), Tim4-PCP5.5 (clone RMT4-54, BioLegend, #130020), CD64-PE/Cy7 (clone X54-5/7.1, BioLegend, #139313), CD11b-PE-Fire640, (clone M1/70, BioLegend, #101279), TCR-b-APC (clone H57-597, BioLegend, #109212), FcERla-A700 (clone MAR-1, BioLegend, #134323), Xcr1-APC/Cy7 (clone ZET, BioLegend, #148223), and CD45-APC-Fire810 (clone 30F11, BioLegend, #103173). Cells were incubated for 30min on ice and washed twice with 1X PBS containing 1% FBS (V/V).

Flow cytometry data was acquired using a Cytek Aurora equipped with violet (405nm), blue (488nm), and red (640nm) lasers. Flow cytometry data was analysed using Flowjo 10 software. Dimensionality reduction and cluster identification was performed using the UMAP and Phenograph packages, respectively. In addition, conventional gating strategies were employed to identify cell populations. Macrophages and cDC2s were determined on their expression of CD64 and MHCII. Subsequently, using antibody labelling-based annotation cDC1s were determined to be MHCII^hi^XCR1^hi^, Langerhans cells assigned as CD207^pos^MHCII^hi^, γδ-T cells as TCRgd^pos^TCRb^neg^, T cells as TCRb^pos^, Neutrophils as CD11b^hi^CCR2^lo^ and Monocytes as CD11b^hi^CCR2^hi^. Mast cells were determined to be FcERla^pos^MHCII^neg^ and B-cells as FcERla^neg^MHCII^hi^. Finally, ILC2s were determined based on Ly6C^pos^CCR2^hi^ expression and NK-cells as Ly6C^pos^CCR2^neg^.

### Organoid cultures

#### Epithelial cell organoid culture

Cells were isolated as described above. After sorting cells were resuspended in organoid medium (ENR) (DMEM/F-12, #11320033, Gibco; 1X PenStrep, #15140122, Gibco; 1X B27 supplement, #17504044, Gibco; 1X GlutaMAX^TM^ supplement, #35050061, Gibco; 0,1% of 100 mM N-Acetyl-cysteine, #A9165-25G, Sigma-Aldrich; 200ng/ml recombinant mouse R-spondin 1, #3474-RS-250, Biotechne; 100ng/ml murine Noggin, #250-38, Peprotech; 50ng/ml murine EGF, #2028-EG-200, R&D systems) and Corning® Matrigel® Growth Factor Reduced (GFR) Basement Membrane Matrix (Matrigel) (Fisher Scientific, #356231) at a 30/70 ratio. 10µl Matrigel domes were plated on preheated 48-well plates (Sarstedt, #83.3923.005) inverted and allowed to dry at 37°C for 20min. Domes were covered in organoid medium (unless stated otherwise) and supplemented with 10µM Y-27632 dihydrochloride (Sigma-Aldrich, #Y0503-5MG) for the first 2 days of culture. Afterwards, medium was changed every second day. For organoid culture treatments, a variety of media was used. ENR^lo^ medium is supplemented with 20ng/ml recombinant R-spondin 1 only (Biotechne, #3474-RS-250). EN^lo^R^lo^ medium contains only 10ng/ml murine Noggin (Peprotech, #250-38) and 20ng/ml recombinant R-spondin 1 (Biotechne, #3474-RS-250). All supplemented media are based on ENR^lo^ medium and contain either, 250ng/ml recombinant IGF1 (Peprotech, #100-11), 250ng/ml recombinant WNT5A (R&D systems, #645-WN-010), 100ng/ml BMP4 (R&D systems, #5020-BP-010), 5ng/ml IL-17A (R&D systems, #421-ML-025/CF), or 100ng/ml recombinant NRG1 (Biotechne, #9875-NR-050).

#### Epithelial and fibroblast co-cultures

Epithelial basal progenitor cells and fibroblasts were isolated as described above and plated at either a 1:1 or 1:2 ratio as stated in the figure legends. The combined cell mixtures were resuspended in minimal organoid medium (DMEM/F-12, Gibco, #11320074; 1X PenStrep, Gibco, #15140122; 1X B27 supplement, Gibco, #17504001; 1X GlutaMAX^TM^ supplement, Gibco, #35050061); 0,1% of 100mM N-Acetyl-cysteine, Sigma-Aldrich, #A9165; 20ng/ml recombinant mouse R-spondin 1, Peprotech, #315-32; 50ng/ml murine EGF, Biotechne, #2028-EG-200) and Matrigel (Fisher Scientific, #356231) at a 30/70 ratio. 10µl domes were plated as above. For the first 2 days of culture, 10µM Y-27632 (Sigma-Aldrich, #Y0503) was added to the medium. Afterwards, the minimal organoid medium was exchanged every second day. Living organoids were imaged using a Leica DMI6000 microscope equipped with a heating chamber and kept at 37°C during the entire acquisition time. Z-stacks were acquired taking an image every 25µm for the entire thickness of the dome.

Organoid size was determined using ImageJ2 (v2.90/1.53t). Organoids encircled at the biggest slice plane using the freehand selection tool. Subsequently, the areas of the encircled section/interface were calculated using the ImageJ *Measure* function.

### Single-cell RNA library preparation

Single-cell experiments were prepared using either Smart-seq3 or Smart-seq3 low volume (Smart-seq3 LV) library preparations as previously described ^21,22^. In brief, for Smart-seq3 library preparations, cells were isolated according to the above described procedure and subsequently sorted into 384 well plates containing 3uL lysis buffer; 0,15% Triton-X100 (Sigma-Aldrich, #93443), 0,4µM/µL RNase Inhibitor (TakaraBio, #2313B), 5% Poly ethylene glycol (PEG) 8000 (Sigma-Aldrich, #P2139) adjusted to reverse transcription (RT) total volume, 0,5µM Smartseq3 OligodT30VN (IDT; 5’-Biotin-ACGAGCATCAGCAGCATACGAT_30_VN-3’) adjusted to RT volume and 0,5mM dNTPs/each (Thermo Fisher, #R0181). After cell sorting lysis plates were centrifuged before storage at -80°C. To process the sorted plates, initial denaturation at 72°C for 10min was performed followed by the addition of 1µL RT mix consisting of 25mM Tris-HCL pH 8.4 (Fisher Scientific, #NC1584154), 30mM NaCl (Ambion, #AM9759), 0.5mM GTP (Thermo Fisher Scientific, #R0461), 2.5mM MgCl2 (Ambion, #AM9530G), 8mM DTT (Thermo Fisher Scientific, #R0861), 0.5U µl^-1^ RNase Inhibitor (Takara Bio, #2313B), 2.0µM Template Switching Oligo (TSO) (5′-Biotin-AGAGACAGATTGCGCAATGNNNNNNNNrGrGrG-3′; IDT) and 2U µl^−1^ of Maxima H Minus reverse transcriptase (Thermo Fisher Scientific, #EP0752). Plates were quickly centrifuged before incubation at 42°C for 90min, followed by ten cycles of 50°C for 2min and 42°C for 2min, and 5min at 85°C. After RT 6µL of PCR was dispensed to each well; 1× KAPA Hotstart HiFI PCR buffer (Roche, #THSNTKB), 0.02 U µl^−1^ of KAPA HiFi Hotstart polymerase (Roche, #THSNTKB), 0.5µM Smartseq3 forward (5′-TCGTCGGCAGCGTCAGATGTGTATAAGAGACAGATTGCGCAATG-3′; IDT), 0,1µM reverse primer (5′-ACGAGCATCAGCAGCATACGA-3′; IDT), 0.5mM MgCl2 (Ambion, #AM9530G) and 0.3mM dNTPs/each (Thermo Fisher Scientific, #R0181). Plates were quickly spun down before being incubated as follows: 3min at 98°C for initial denaturation, 22 cycles of 20 seconds at 98°C, 30 seconds at 65°C and 4min at 72°C. Final elongation was performed for 5min at 72°C. After PCR pre-amplified cDNA were clean-up using 22% home-made PEG beads at a ratio of 0.7 beads:1 sample before concentration quantification using QuantiFluor® dsDNA System (Promega, #E2671). cDNA was normalised and diluted to 100pg µl^-1^.

For Smart-seq3 LV library preparation each well was dispensed with 3µL of Vapor-lock (Qiagen) prior addition of lysis buffer. All steps and reagent concentrations are like the Smart-seq3 procedure above, except all reaction volumes have been scaled 1:10, and are carried out underneath the hydrophobic overlay Vapor-lock (QIAGEN, #981611). Instead of cleaning up the cDNA, pre-amplified cDNA libraries were diluted with addition of 9µL water to each well before quantification using QuantiFluor^®^ dsDNA System (Promega, #E2671) and normalisation and dilution to 100pg ul^-1^.

Tagmentation was performed similar for both Smart-seq3 and Smart-seq3 LV in 2µL volume, consisting of 1µL normalised cDNA (100 pg) and 1X tagmentation buffer (10mM Tris pH 7.5, 5mM MgCl2, 5% DMF, Thermo Fisher Scientific, #20673), 0.5µl of ATM (Illumina XT DNA sample preparation kit, #FC-131-1096). Plates were incubated for 10min at 55°C before the reaction was stopped by the addition of 0.5µL 0.2% SDS (Thermo Fisher Scientific, #24730020) to each well. Index PCR was carried out after the addition of 1.5µL custom Nextera Index primers (0.5µM) by dispensing 4µL of PCR mix; 1× Phusion Buffer (Thermo Fisher Scientific, #F631L), 0.01 U µl^−1^ of Phusion DNA polymerase (Thermo Fisher Scientific, #F630L), 0.2mM dNTP each (Thermo Fisher Scientific, #R0181). PCR was performed out at 3min at 72°C; 30 seconds at 95°C; 12 cycles of (10 seconds at 95°C; 30 seconds at 55°C; 30– 60 seconds at 72°C); and 5min at 72°C. Each final and indexed library plate was spun out gently using a 300-ml robotic reservoir (Nalgene) fitted with a custom 3D-printed scaffold by pulsing the centrifuge to < 200*g.* The pooled libraries were afterwards purified with home-made 22% PEG beads at a ratio of 1 sample to 0.7 beads.

### Sequencing

Final Smart-seq3 and Smart-seq3 LV libraries were sequenced on a MGI DNBSEQ G400RS platform (version 1.1.0.108 software). To prepare for sequencing on the MGI platform, circular ssDNA libraries were created using the MGIEasy Universal Library Conversion Kit (MGI). Adapter conversion PCR was carried out on 50ng of final pooled library for 5 cycles, following circularisation of 1pmol dsDNA according to manufacturer’s protocol. DNA nanoballs (DNBs) were created from 60fmol of circular ssDNA library pools using a custom rolling-circle amplification primer (5′-TCGCCGTATCATTCAAGCAGAAGACG-3′, IDT). DNBs were sequenced PE100 using the following custom sequencing primers

Read 1: 5′-TCGTCGGCAGCGTCAGATGTGTATAAGAGACAG-3′;

MDA: 5′-CGTATGCCGTCTTCTGCTTGAATGATACGGCGAC-3′,

Read 2: 5′-GTCTCGTGGGCTCGGAGATGTGTATAAGAGACAG-3′;

i7 index: 5′-CCGTATCATTCAAGCAGAAGACGGCATACGAGAT-3′; and

i5 index: 5′-CTGTCTCTTATACACATCTGACGCTGCCGACGA-3′ on a FCL flow cell.

### Primary single-cell data processing

zUMI version 2.9.7 was utilised to process raw FASTQ files ^62^. UMI-containing reads in case of Smart-seq3xpress were identified by the (ATTGCGCAATG) pattern allowing up to two mismatches. Low-quality barcodes and UMIs were used to filter reads (4 bases < phred 20, 3 bases < phred 20, respectively). Reads were mapped to the mouse genome (GRCm39/mm39) using STAR version 2.7.3a. Read counts and umicounts were calculated using Gencode GRCm39 vM29 annotation.

### Single-cell analysis

Cells were filtered for low-quality libraries requiring the following criteria; more than 40 % read-pairs mapped to exons, more than 10.000 read-pairs sequenced, more than 500 genes (exon+intron quantification) detected per cell, more than 1000 UMIs (exon+intron quantification) detected per cell, and less than 25% of read pairs mapped to mitochondrial genes. Furthermore, a gene was required to be expressed in at least ten cells. Analysis was performed using Seurat (v4.3.0) ^63^. Data was normalised, scaled to 10,000, and total number of counts and the mitochondrial gene fraction was regressed out. Using the integration function, the effects from our 3 different cell isolations (“Batch”) were except for Fibroblasts run1 and other cells run1 and run2 due to very low cell numbers. The top 3,000 variable genes were considered and 50 principal components for shared nearest neighbour (SNN) neighbourhood construction and UMAP dimensionality reduction. Cell clusters were produced using Louvain algorithm at a resolution of 0.8. Differential gene expression across cell identities was determined utilising the Seurat function *FindAllMarkers* (with default parameters except *logfc.threshold = 0.58 to enrich for genes that are at least 1.5 fold changed*).

#### Subclustering

Subclustering of epithelial cells, fibroblasts, immune cells, and all other cell types was performed based on the obtained Louvain algorithm clusters and annotation was conducted based on known marker expression. Epithelial cells were grouped based on expression of *Krt5, Krt14, Krt4, Krt13*, and *Krt8, and Krt18* expression was used to exclude stomach epithelial cells. Fibroblasts were identified based on expression of *Pdgfra*. Immune cells based on *Ptprc* expression and known CD45 fluorescence intensity, endothelial cells based on *Pecam1* expression, enteric nerve cells based on expression of *Sox10*, Pericytes based on expression of *Pdgfra* and *Pdgfrb*. Lymphatic vessels based on *Pecam1* and *Lyve1*, smooth muscle cells on *Acta2* and *Actg2*, and muscle cells based on *Pax7*. For each subclustered major cell type the *FindAllMarkers* function with default parameters was run to determine differentially expressed genes. The obtained DEGs were used for cell identity annotation (Supplemental Table 12).

#### Identification of regional differentially expressed genes

To identify regional differentially expressed genes each subclustered major cell type identity was changed to the information contained in the *Region2* column of the metadata file. Seurat *FindMarkers* function was run with default parameters (except *logfc.threshold = 0, min.pct.diff = 0*) setting *ident.1 = “Proximal”* and *ident.2 = “Distal”*. Determination of DEGs between epithelial basal cell populations was performed using Seurat *FindMarkers* with default parameters setting *ident.1 = “Basal 3”* and *ident.2 = c(“Basal 1”, “Basal 3”, “Basal 4”, “Basal 5”)*. Gene scoring analysis for epithelial population *Basal 3* was performed using the Seurat *AddModuleScore* function with *Lrig1*, *Pappa*, *Igfbp5*, and *Smoc2* as the gene list.

#### Systematic inference of cell-cell communication

CellChat (v1.5) was used to determine cell-cell communication patterns using the ligand-receptor interaction database CellChatDB. CellChat infers cell-cell communication based on over-expressed ligands and receptors for each cell group. Average expression of a ligand in one cell group and the corresponding receptor in another cell group as well as the expression of cofactors is then used to calculate communication probabilities between cell groups. Significant interactions are determined utilising permutation tests. The calculated communication probabilities of all associated ligand-receptor pairs within a given signalling pathway are then summarised to generate an intercellular communication network. We utilised CellChat to calculate inferred cell-cell communication on the subclustered and merged Seurat objects of epithelial cells and fibroblasts with regional identities.

#### Cell-cell communication across oesophageal regions

Moreover, we wanted to investigate signalling changes across regions. In a first step, we merged Seurat objects of subclustered regionalized epithelial cells and fibroblasts. Comparing the obtained proximal and distal dataset we compare the number of cell-cell interactions across the two regions. Secondly, we merged Seurat objects of all subclustered major cell types from the proximal or distal oesophagus, respectively. The generated proximal and distal datasets were merged in CellChat and subjected to overall comparison of signalling pathways and altered ligand-receptor pairs between the proximal and distal oesophagus. Our aim was to identify cell populations with significant interaction changes along the oesophageal length as well as to identify specific regional ligand-receptor pairs between cell groups. We use various built in CellChat visualization functions to present our analysis.

### Statistical analyses

GraphPad Prism (v 9.5.1) and R (v 4.1.1) have been used for statistical analyses. Statistics were assessed by two-tailed unpaired or ratio-paired t-test, or ordinary one-way analysis of variance (ANOVA) as indicated in the figure legends. ANOVA-based analysis was followed by a Holm-Šídák’s multiple comparisons test for multiple comparisons. Error bars represent the mean SD.

## Acknowledgements

This study was supported by ERC Starting Grant (TroyCAN 851241), Cancerfonden (19 0007Pj), SFO StratRegen 2023/24 Junior Grant and Vetenskapsrådet (2023-02743). MG is a Cancerfonden Senior Investigator. We are grateful for all technical assistance from Karolinska Institutet core facilities; especially Florian Salomons of the Biomedicum Imaging Core. We thank Rebecca Cardoso and Eduardo Villablanca for the IL-17A reporter mouse. We thank Melanie Pieber, Jinming Han, Monika Plescher, and Jannis Kalkitsas for help with animal experiments and the Genander lab for constructive feedback on the manuscript. Parts of the figures were drawn by using Servier Medical Art which is licensed under a Creative Commons Attribution 3.0 Unported License (https://creativecommons.org/licenses/by/3.0/).

## Author contributions

**DG**: Conceptualization, Methodology, Software, Validation, Formal analysis, Investigation, Data Curation, Writing – original draft preparation, Writing – review and editing, Visualization, Project administration, Supervision **MH-J**: Conceptualization, Methodology, Software, Validation, Formal analysis, Investigation, Data Curation, Writing – review and editing, Supervision **HL**: Methodology, Validation, Formal analysis, Investigation, Visualization **EE**: Validation, Investigation, Writing – review and editing **AJ**: Investigation **CG**: Resources **RAH**: Resources **RS**: Resources, Funding acquisition **MG**: Writing – original draft preparation, Writing – review and editing, Supervision, Resources, Funding acquisition. All authors have proof-read and approved the final version of the manuscript.

---

**Supplemental Figure 1. Related to Figure 1.**
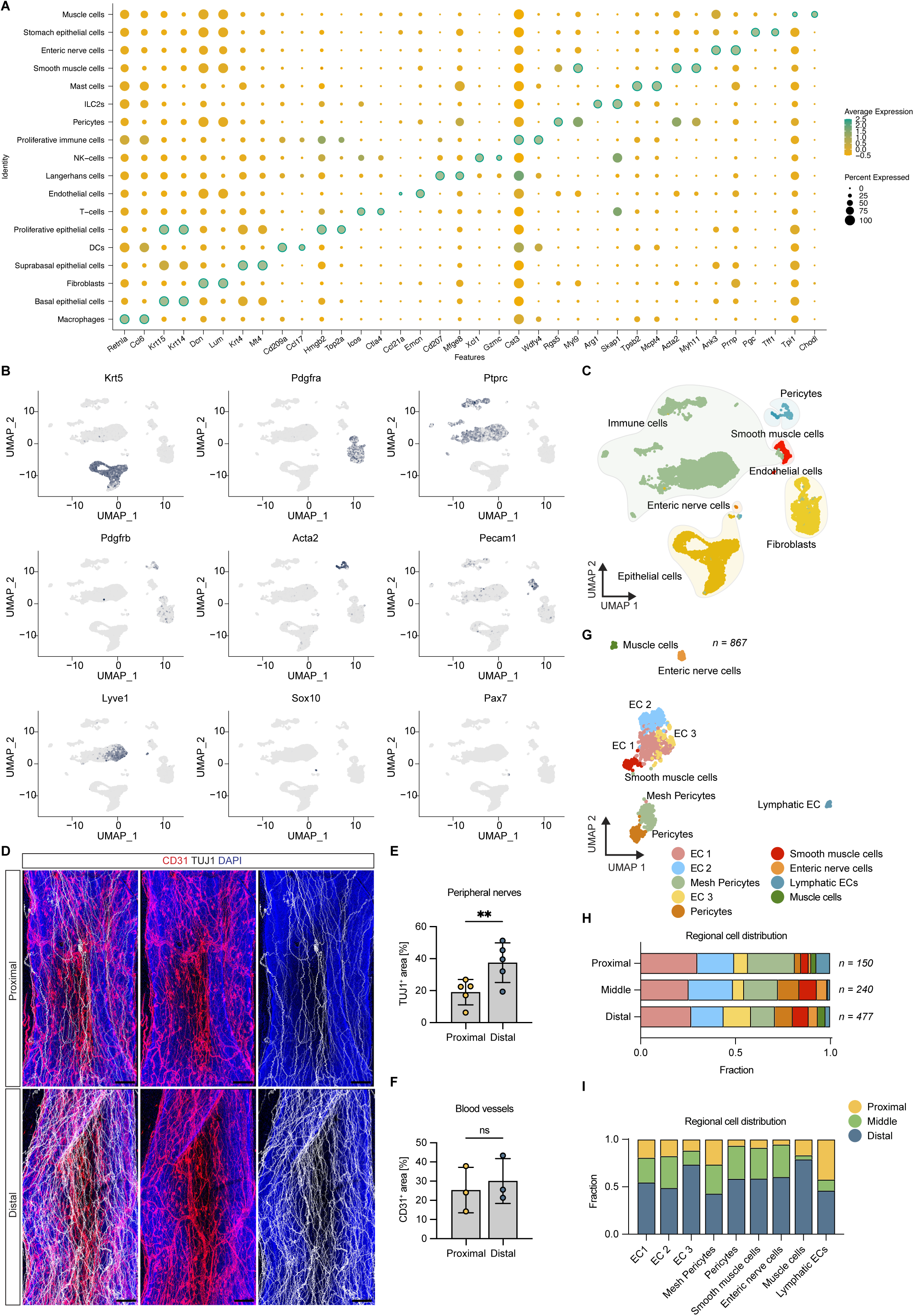
(A) DotPlot of the two top markers for each cell type and state annotated based on the entire data set. (B, C) UMAP visualizations of markers (B) used to group cell types (epithelial cells, fibroblasts, immune cells, pericytes, smooth muscle cells, endothelial cells and peripheral nerves) displayed in (C). (D) Maximum intensity projections of proximal and distal oesophageal whole mount stainings, using CD31 (red) and TUJ1 (white) to visualize vessels and nerve fibres respectively. Scale bar = 200μm. (E) Quantification of TUJ1+ positive area on proximal and distal whole mounts. n = 5 mice. (F) Quantification of CD31+ positive area on proximal and distal whole mounts. n = 3 mice. (G) UMAP of individually subclustered stromal cell types other than fibroblasts and immune cells. EC1-3 are endothelial cell subpopulations. (H) Quantification of the distribution of stromal cells within each sequenced proximal-distal region. Cell numbers were normalised within the respective region and resulting fractions are displayed. (I) Relative stromal cell distribution within each oesophageal region. (E, F) Ratio paired two-sided t-test. ns p > 0.05, * p < 0.05, ** p < 0.01, *** p < 0.001.

**Supplemental Figure 2. Related to Figure 2.**
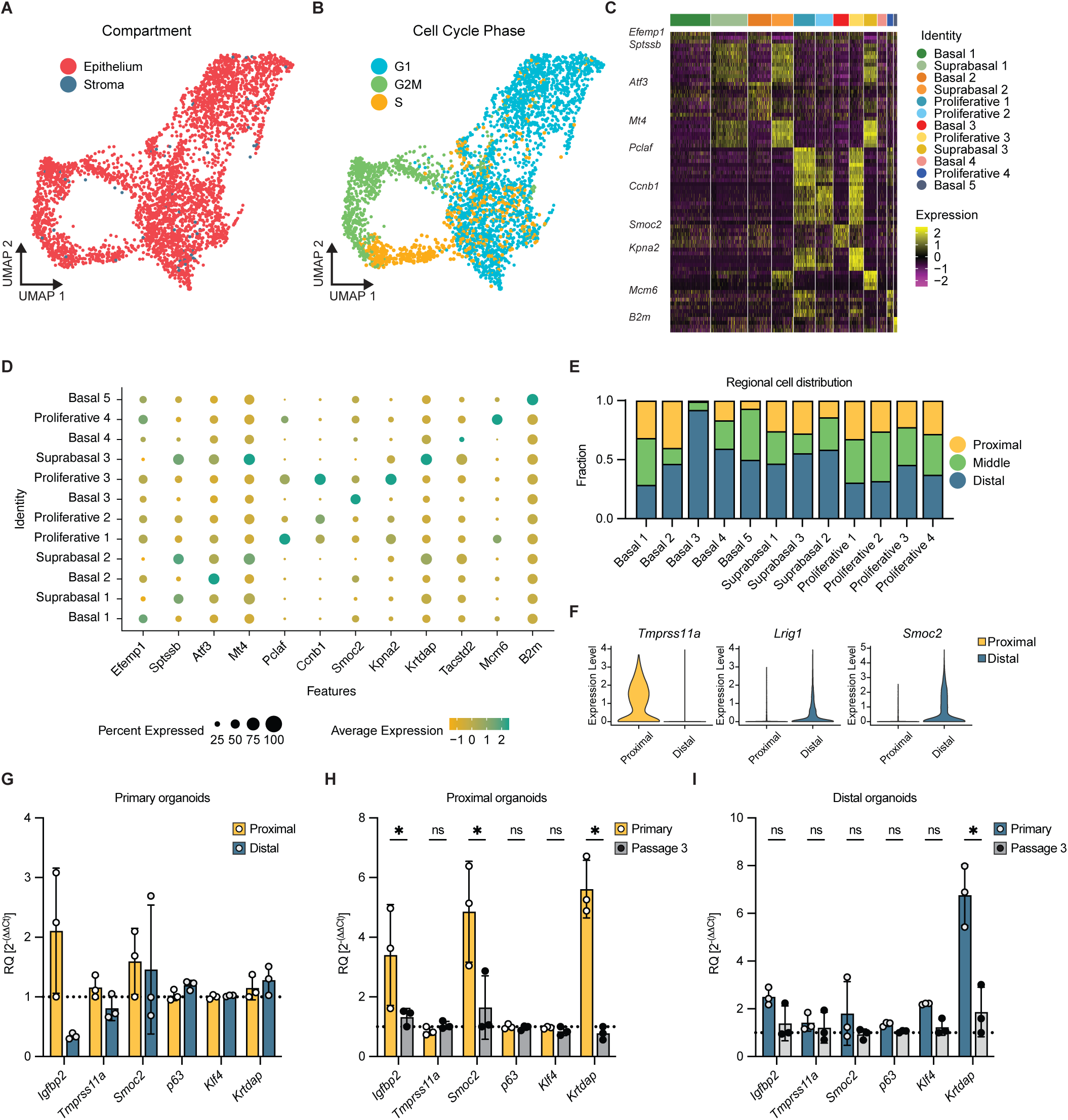
(A) UMAP of subclustered epithelial cells colour-coded by oesophageal compartment-of- origin. (B) UMAP of all subclustered epithelial cells colour-coded by inferred cell-cycle stage. (C) Heatmap of top 10 DEGs of subclustered epithelial subpopulations. (D) DotPlot of top differentially expressed gene (log2 fold-change) of all epithelial subpopulations. (E) Distribution of each epithelial subpopulation across oesophageal regions, normalised within cell identity. (F) Violin plots showcasing differentially expressed *Lrig1*, *Tmprss11a* and *Smoc2* comparing all cells in the proximal and distal epithelium. (G) Gene expression of proximal and distal primary organoid cultures grown in ENRlo medium. (H) Gene expression comparing proximal primary to passaged organoids. (I) Gene expression comparing distal primary to passaged organoids. Regional signature profiles are lost or altered upon culture. n = 3 (2-3 mice per n). Figure (G-I) Multiple ratio paired t-test corrected for multiple comparisons with Holm- Šídák method. ns p > 0.05, * p < 0.05, ** p < 0.01, *** p < 0.001.

**Supplemental Figure 3. Related to Figure 4.**
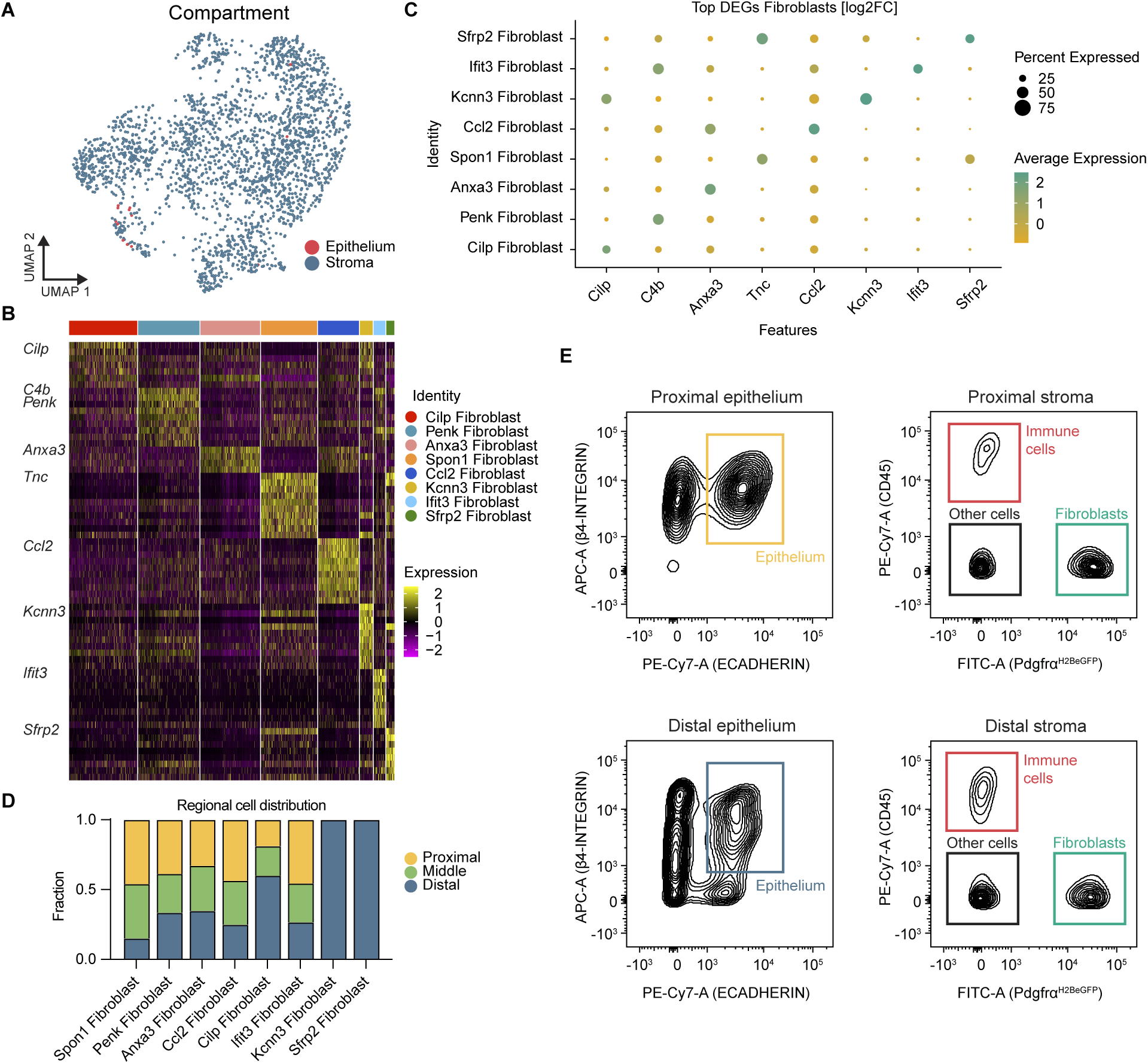
(A) UMAP of subclustered fibroblasts colour-coded by oesophageal compartment. (B) Heatmap of top 10 DEGs of fibroblasts subpopulations displayed in Figure 4A. (C) DotPlot of the top differentially expressed gene (log2 fold-change) for each fibroblast subpopulation. (D) Distribution of fibroblast subpopulations across oesophageal regions, normalised within cell identity. (E) FACS isolation strategy for proximal and distal ECAD^pos^/ITGB4^pos^ epithelial (left panels) and Pdgfrα:H2BeGFP^pos^,CD45^neg^ stromal (right panel) cells used for organoid co- cultures.

**Supplemental Figure 4. Related to Figure 5.**
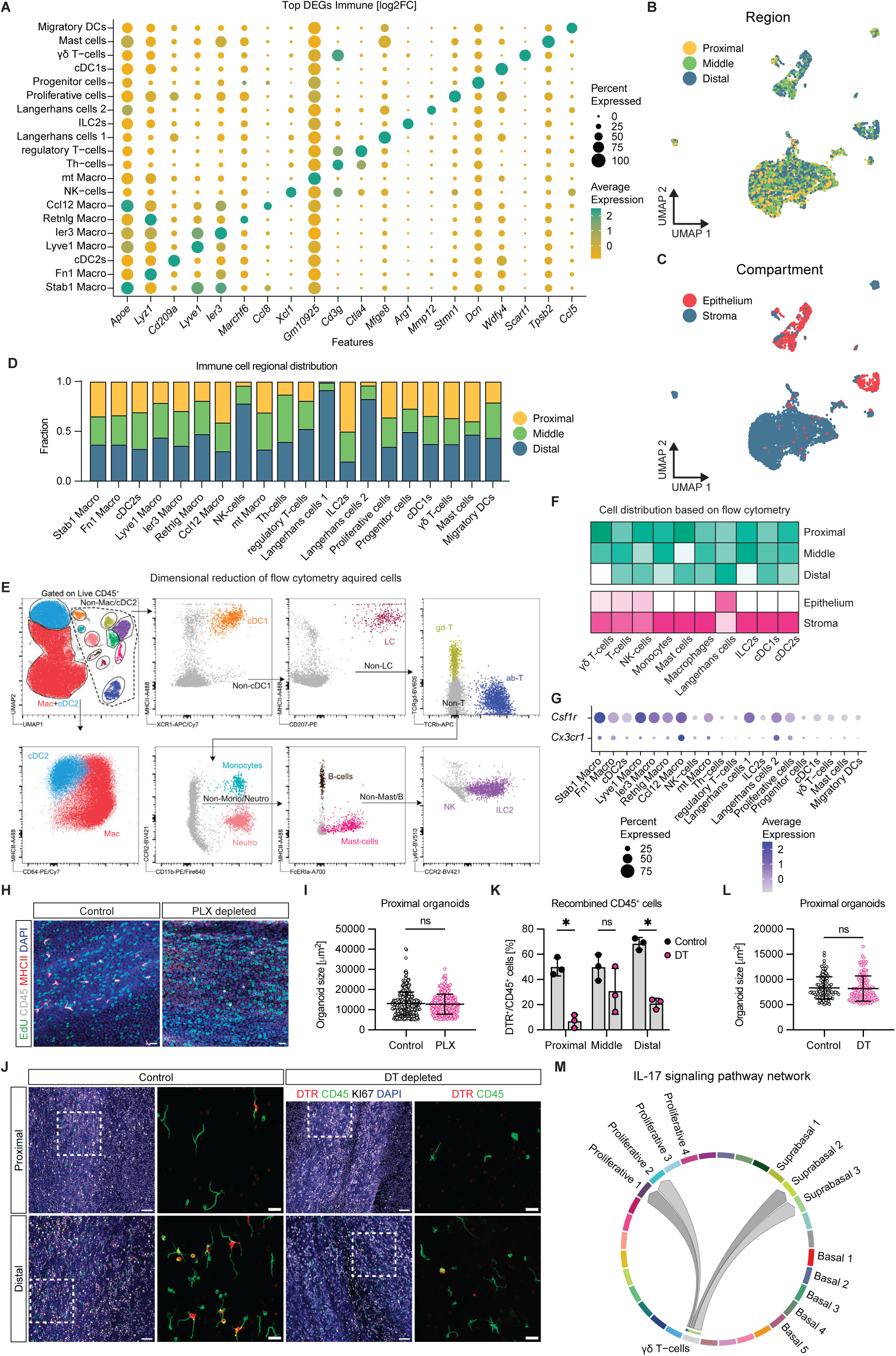
(A) DotPlot of the top differentially expressed gene (log2 fold-change) of subclustered immune cell types. (B) UMAP of subclustered immune cells colour-coded by oesophageal region. (C) UMAP of subclustered immune cells colour-coded by oesophageal compartment. (D) Distribution of subclustered immune cells across oesophageal regions normalised within cell identity. (E) UMAP dimensional reduction plot and gating strategy for flow cytometry determined immune cell identities. (F) Heatmaps displaying the relative regional (green) and compartmental (pick) distribution of spectral flow cytometry analysed immune cell identities. (G) *Csf1r* and *Cx3cr1* expression in immune cells. (H) Representative immunofluorescent images of epithelial whole mounts of control and PLX3397 chow mice counterstained for EdU (green), CD45 (white), MHCII (red), and DAPI (blue). Scale bar = 20µm. (I) Organoid size of proximally derived organoids from control and PLX3397 chow mice. n = 3 (2 mice per n). Each dot represents one organoid. (J) Representative immunofluorescent images of oesophageal epithelial whole mounts of control mice and Cx3cr1-CreER:DTR/DTR immune cell depleted mice counterstained for KI67 (white), CD45 (green), DTR (red), and DAPI (blue). Scale bar = 50 µm and inlets 20 µm. (K) Quantification of DTR-positive, recombined, CD45- positive cells in the proximal, middle and distal oesophagus comparing control (no DT) to DT injected mice. (L) Comparing size of organoids derived from proximal basal cells in control and DT treated mice. n = 3 with 2 mice per n. Each dot represents one organoid. (M) Cellchat- inferred probable communication, indicating that IL-17A, produced by intra-epithelial ψ8T- cells, impacts proliferating as well as differentiating epithelial cell populations. (I, L) Two- tailed Kolmogorov-Smirnov test. ns p > 0.05, * p < 0.05, ** p < 0.01, *** p < 0.001. (K) Multiple unpaired t-test corrected for multiple comparisons with Holm-Šídák method. ns p > 0.05, * p < 0.05, ** p < 0.01, *** p < 0.001.

**Supplemental Figure 5. Related to Figure 6.**
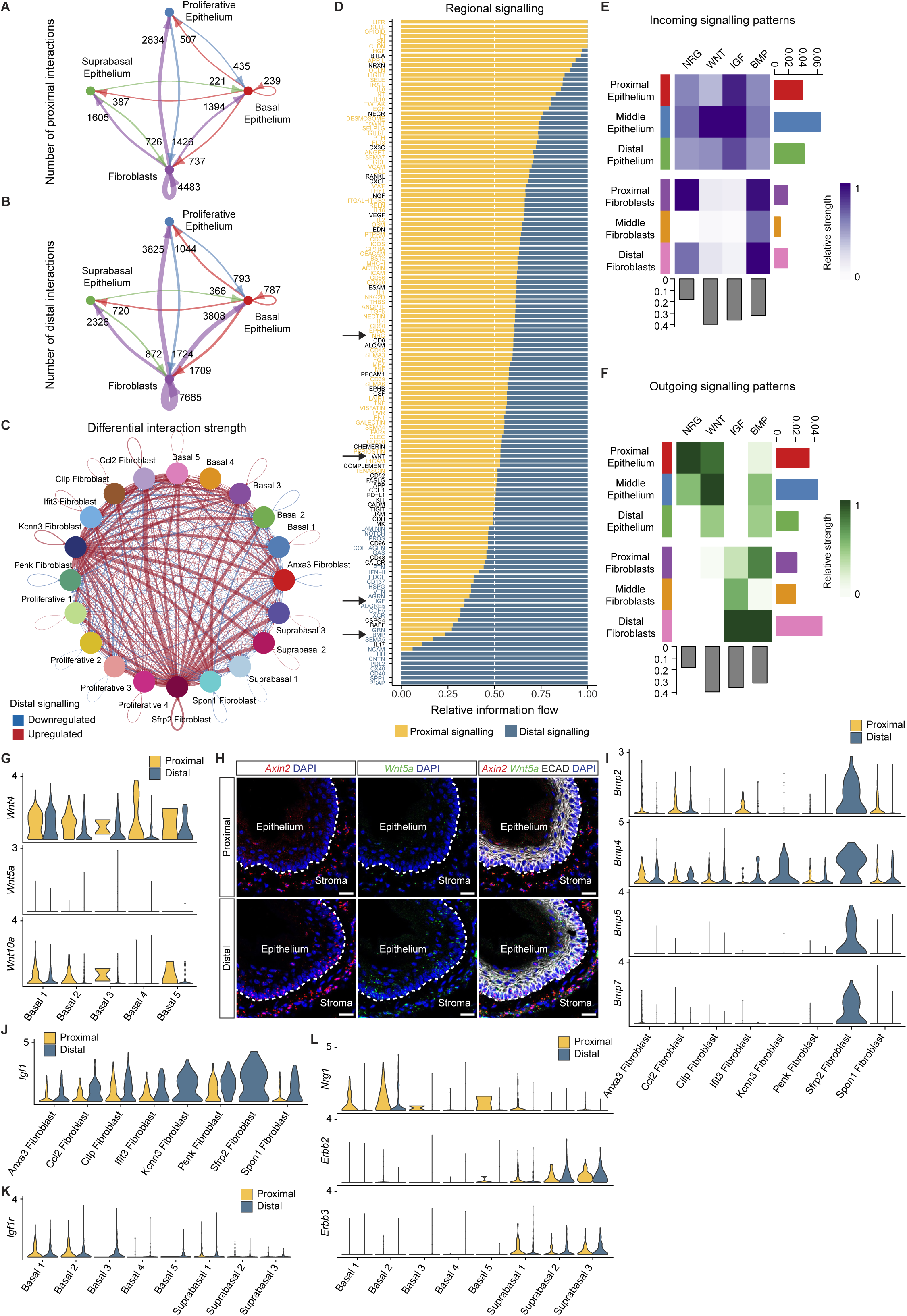
(A-B) Circular plot displaying number of interactions between proximal fibroblasts and proximal epithelial cells (A) and between distal fibroblasts and distal epithelial cells (B). Epithelial cells were aggregated into basal, suprabasal, and proliferative cells based on the annotation of subclustered epithelial cells. (C) Circular plot of differential interaction strengths in cell-cell communication between fibroblasts and epithelial cells comparing datasets containing proximal epithelial cells and fibroblasts and distal epithelial cells and fibroblasts, respectively. (D) Information flow (sum of communication probabilities among all pairs of cell groups) of all significantly altered signalling pathways comparing proximal and distal derived cells. (E, F) Heatmaps of selected incoming (blue) (E) and outgoing (green) (F) signalling pathways of regional oesophageal fibroblasts and epithelial cells. (G) Violin plots displaying regional gene expression of expressed WNT ligands in epithelial basal cells. (H) *In situ* hybridization of *Axin2* (red) and *Wnt5a* (green) on cross-sections of the proximal and distal oesophagus. Sections are counterstained with E-CADHERIN (ECAD, white) and DAPI (blue). Scale bar = 20 µm. Dotted line marks the epithelial-stromal border. (I-J) Violin plots displaying regional gene expression of selected BMP ligand (I) and *Igf1* (J) in fibroblasts. (K-L) Violin plot displaying regional gene expression of *Igf1r* (K) and *Nrg1* (L) in basal and suprabasal epithelial cells.

**Supplemental Figure 6. Related to Figure 7.**
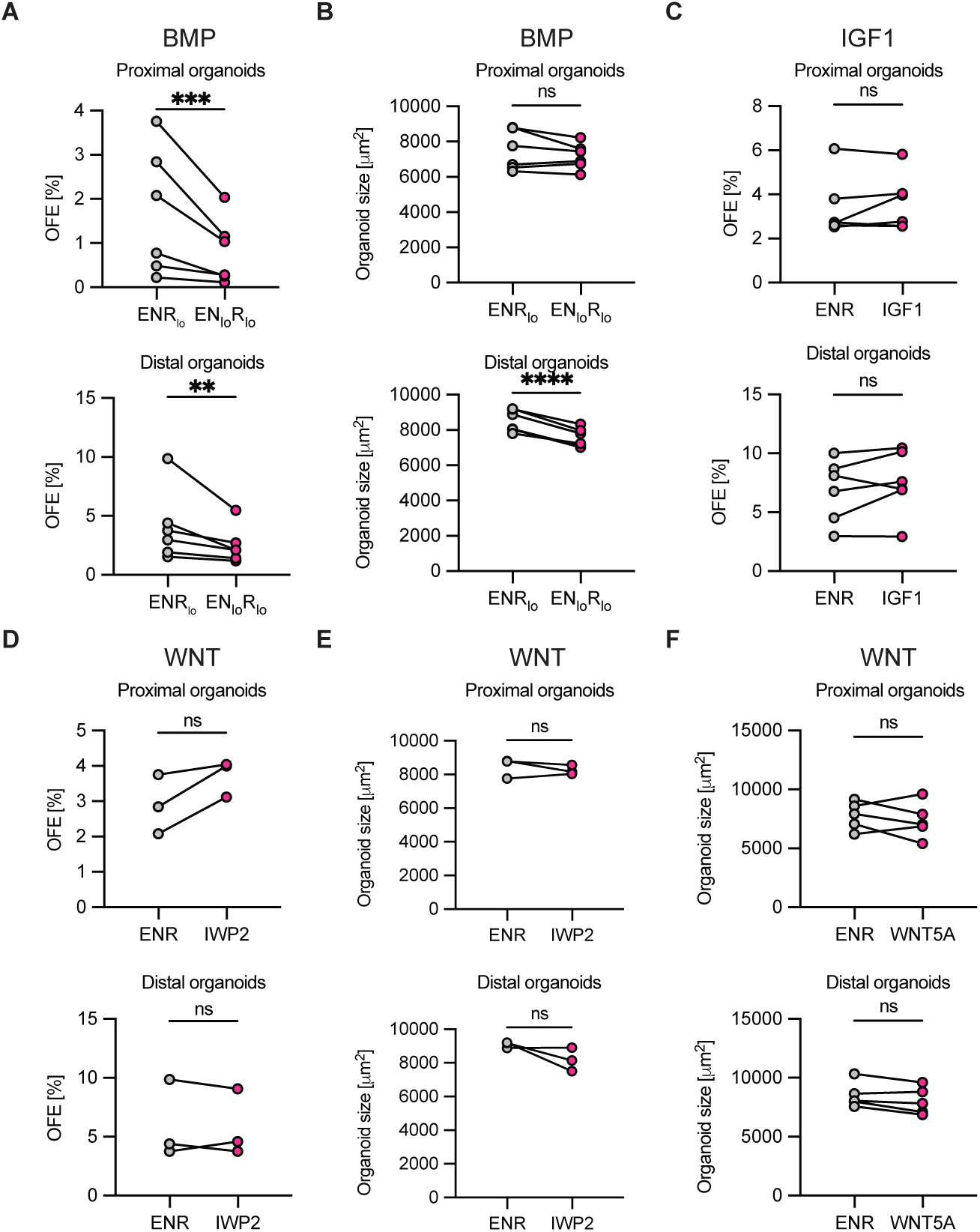
(A-B) Quantification of proximal and distal organoid forming efficiency (A) and size (B) in control medium and medium with reduced NOGGIN (10ng/mL). (C) Quantification of proximal and distal organoid forming efficiency in control medium and medium supplemented with 250ng/mL IGF1. (D-E) Quantification of proximal and distal organoid forming efficiency (D) and size (E) in control and IWP2 (5µM) containing medium, inhibiting WNT secretion. (F) Quantification of proximal and distal organoid size in control medium and medium supplemented with 250ng/mL WNT5A. Each dot represents OFE or the average size of all organoids. n=3-6. (M, N, O, P) Two-sided ratio paired t-test. ns p > 0.05, * p < 0.05, ** p < 0.01, *** p < 0.001.

**Supplemental Table 1. Differentially expressed genes in stromal cells.**

Table of DEGs for clusters among niche cells displaying p-value, average log2 fold-change, percentage of expression within cluster (pct.1), percentage of expression in all other clusters (pct.2), and adjusted p-value.

**Supplemental Table 2. Gene ontology analysis of stromal cells.**

Table of identified GO terms displaying GO term ID, description, gene ratio, background ratio, p-value, adjusted p-value, q-value, gene IDs, and gene count for each identified GO term for each niche cell cluster separated in sheets.

**Supplemental Table 3. Differentially expressed genes in the epithelium.**

Table of DEGs for clusters among epithelial cells displaying p-value, average log2 fold- change, percentage of expression within cluster (pct.1), percentage of expression in all other clusters (pct.2), and adjusted p-value.

**Supplemental Table 4. Gene ontology analysis of the epithelium.**

Table of identified GO terms displaying GO term ID, description, gene ratio, background ratio, p-value, adjusted p-value, q-value, gene IDs, and gene count for each identified GO term for each epithelial cell cluster separated in sheets.

**Supplemental Table 5. Regional DEGs in the epithelium.**

Table of DEGs identified comparing all gene expression of all proximal epithelial cells to all distal epithelial cells. Displayed are p-value, average log2 fold-change, percentage of expressing cells proximally (pct.1), percentage of expressing cells distally (pct.2), and adjusted p-value for each gene.

**Supplemental Table 6. DEGs of epithelial Basal 3 cluster.**

Table of DEGs identified comparing epithelial subcluster ‘Basal 3’ against the other four epithelial basal cell subclusters. Displayed are p-value, average log2 fold-change, percentage of expressing cells Basal 3 (pct.1), percentage of expressing cells in other basal cell clusters (pct.2), and adjusted p-value for each gene.

**Supplemental Table 7. Differentially expressed genes in fibroblasts.**

Table of DEGs for clusters among fibroblasts displaying p-value, average log2 fold-change, percentage of expression within cluster (pct.1), percentage of expression in all other clusters (pct.2), and adjusted p-value.

**Supplemental Table 8. Gene ontology analysis of fibroblasts.**

Table of identified GO terms displaying GO term ID, description, gene ratio, background ratio, p-value, adjusted p-value, q-value, gene IDs, and gene count for each identified GO term for each fibroblast cluster separated in sheets.

**Supplemental Table 9. Regional DEGs of fibroblasts.**

Table of DEGs identified comparing all gene expression of all proximal fibroblasts to all distal fibroblasts. Displayed are p-value, average log2 fold-change, percentage of expressing cells proximally (pct.1), percentage of expressing cells distally (pct.2), and adjusted p-value for each gene.

**Supplemental Table 10. Differentially expressed genes in immune cells.**

Table of DEGs for clusters among immune cells displaying p-value, average log2 fold-change, percentage of expression within cluster (pct.1), percentage of expression in all other clusters (pct.2), and adjusted p-value.

**Supplemental Table 11. Gene ontology analysis of immune cells.**

Table of identified GO terms displaying GO term ID, description, gene ratio, background ratio, p-value, adjusted p-value, q-value, gene IDs, and gene count for each identified GO term for each immune cell cluster separated in sheets.

**Supplemental Table 12. Subclustered cell numbers per region.**

Table of all subclustered cell types containing cell numbers per region and identity.

